# ZHX2 Promotes HIF1α Oncogenic Signaling in Triple-Negative Breast Cancer

**DOI:** 10.1101/2021.05.27.445959

**Authors:** Wentong Fang, Chengheng Liao, Rachel Shi, Jeremy M. Simon, Travis S. Ptacek, Giada Zurlo, Youqiong Ye, Leng Han, Cheng Fan, Christopher Llynard Ortiz, Hong-Rui Lin, Ujjawal Manocha, Weibo Luo, William Y. Kim, Lee-Wei Yang, Qing Zhang

**Affiliations:** Department of Pharmacy, The First Affiliated Hospital of Nanjing Medical University, Nanjing, Jiangsu 210029, China; Lineberger Comprehensive Cancer Center, University of North Carolina School of Medicine, Chapel Hill, NC 27599, USA; Department of Pathology, University of Texas Southwestern Medical Center, Dallas, TX 75390, USA; Department of Genetics, Neuroscience Center, University of North Carolina, Chapel Hill, NC 27599, USA; UNC Neuroscience Center, Carolina Institute for Developmental Disabilities, University of North Carolina, Chapel Hill, NC 27599, USA; Shanghai Institute of Immunology, Faculty of Basic Medicine, Shanghai Jiao Tong University School of Medicine, Shanghai, 200025, China; Department of Biochemistry and Molecular Biology, The University of Texas Health Science Center at Houston McGovern Medical School, Houston, TX, 77030, USA; Institute of Bioinformatics and Structural Biology, National Tsing Hua University, Hsinchu 300, Taiwan; Chemical Biology and Molecular Biophysics Program, Taiwan International Graduate Program, Institute of Chemistry, Academia Sinica, Taipei 115, Taiwan; Department of Chemistry, National Tsing-Hua University, Hsinchu 30013, Taiwan; Physics Division, National Center for Theoretical Sciences, Hsinchu 30013, Taiwan

**Keywords:** ZHX2, TNBC, HIF1α, VHL

## Abstract

Triple-negative breast cancer (TNBC) is an aggressive and highly lethal disease, which warrants the critical need to identify new therapeutic targets. We show that Zinc Fingers And Homeoboxes 2 (ZHX2) is amplified or overexpressed in TNBC cell lines and patients. Functionally, depletion of ZHX2 inhibited TNBC cell growth and invasion in vitro, orthotopic tumor growth and spontaneous lung metastasis in vivo. Mechanistically, ZHX2 bound with hypoxia inducible factor (HIF) family members and positively regulated HIF1α activity in TNBC. Integrated ChIP-Seq and gene expression profiling demonstrated that ZHX2 co-occupied with HIF1α on transcriptionally active promoters marked by H3K4me3 and H3K27ac, thereby promoting gene expression. Furthermore, multiple residues (R491, R581 and R674) on ZHX2 are important in regulating its phenotype, which correspond with their roles on controlling HIF1α activity in TNBC cells. These studies establish that ZHX2 activates oncogenic HIF1α signaling, therefore serving as a potential therapeutic target for TNBC.

## Introduction

Triple-negative breast cancer (TNBC) accounts for 15-20% of all breast cancer (Anders & Carey, 2009). TNBC is associated with a more aggressive clinical history, a higher likelihood of distant metastasis, shorter survival and a higher mortality rate compared to other subtypes of breast cancer (Anders & Carey, 2009). In addition, recent studies illustrate high rates of brain metastasis in TNBC that is associated with poor survival (Heitz *et al*, 2009; Lin *et al*, 2008; Niwinska *et al*, 2010). Since TNBCs do not express estrogen receptor (ER), progesterone receptor (PR) or human epidermal growth factor receptor 2 (HER2), treatment options have historically been limited to chemotherapy(Masui *et al*, 2013), which has significant toxicity and a suboptimal impact on the five-year relapse rate. Therefore, it is critical to identify novel therapeutic targets in TNBC.

The Zinc-fingers and homeoboxes (ZHX) family includes ZHX1, 2 and 3. ZHX2 is located on 8q24.13 and contains two zinc finger domains and five homeodomains (HDs) (Kawata *et al*, 2003). In addition, between amino acids 408-488, it contains a proline rich region (PRR). ZHX2 can form homodimers or can hetero-dimerize with the other two family members ZHX1 or ZHX3 (Kawata *et al*., 2003). Originally, ZHX2 was found to be a key transcriptional repressor for the alpha-fetoprotein regulator 1 (*Afr1*) (Perincheri *et al*, 2005), which is an important oncogene in liver cancer. From this perspective, ZHX2 was identified and reported to function as a transcriptional repressor (Kawata *et al*., 2003), where fusion of ZHX2 with a GAL4-DNA binding domain repressed transcription of a GAL4-dependent luciferase reporter. Additionally, ZHX2 was reported to have tumor suppressor activity in hepatocellular carcinoma (HCC), by repressing cyclin A, E and multidrug resistance 1 (MDR1) expression (Ma *et al*, 2015; Yue *et al*, 2012). ZHX2 was also indicated to be a tumor suppressor in Hodgkin lymphoma or myeloma although there is no direct experimental evidence supporting this hypothesis (Armellini *et al*, 2008; Nagel *et al*, 2012; Nagel *et al*, 2011).

However, accumulating evidence suggests that ZHX2 may contribute to cancer pathology in other contexts. Tissue microarray and clinicopathological analysis show that ZHX2 protein expression in metastatic HCC is twice as high as in the primary lesions, indicating that ZHX2 expression is associated with metastasis in HCC (Hu *et al*, 2007). In addition, our recent findings through genome-wide screening identify that ZHX2 is a substrate of von Hippel Lindau (gene name *VHL,* protein name pVHL) protein, accumulates in kidney cancer, and promotes oncogenic signaling by at least partially activating NF-κB signaling in clear cell renal cell carcinoma (ccRCC) (Zhang *et al*, 2018). These pieces of evidence suggest that ZHX2 acts as a tumor suppressor or oncogene in a context-dependent manner. It is also important to point out that ZHX2 is located on 8q24, a genomic region that is frequently amplified in various cancers including breast cancer (Guan *et al*, 2007). More importantly, the role of ZHX2 in other cancers, such as in TNBC, remains largely unknown.

Tumor hypoxia is a characteristic of most solid tumors. Hypoxic cells are known to confer radio- or chemotherapeutic resistance, and therefore are hypothesized to undergo positive selection during cancer development (Brown & Wilson, 2004; Gray *et al*, 1953). The key proteins mediating oxygen sensing in these cells involve two classes of proteins: (1) upstream oxygen sensors, namely the prolyl hydroxylases EglN1-3, responsible for the hydroxylation of various substrates, such as hypoxia inducible factor (HIF), FOXO3a, ADSL, SFMBT1 and TBK1 (Hu *et al*, 2020; Liu *et al*, 2020; Semenza, 2012; Zheng *et al*, 2014; Zurlo *et al*, 2019); (2) the downstream VHL E3 ligase complex. For example, EglN family members (EglN1, primarily *in vivo*) hydroxylate HIF1α on proline 402 and 564 positions, which lead to pVHL binding and HIF1α ubiquitination and degradation (Appelhoff *et al*, 2004; Ivan *et al*, 2001; Jaakkola *et al*, 2001). HIF1α has been well established to be an important oncogene in multiple cancers, including breast cancer (Briggs *et al*, 2016; Semenza, 2010). Tumor hypoxia or pVHL loss will lead to the accumulation of HIF1α. As a result of the accumulation and translocation of HIFα factors into the nucleus, HIF1α dimerizes with a constitutively expressed HIF1β subunit (also called ARNT) and transactivates genes that have hypoxia response elements (NCGTG) in promoters or enhancer regions. HIF1-transactivated genes include those involved in angiogenesis (e.g. *VEGF*), glycolysis and glucose transport (e.g. *GLUT1*), and erythropoiesis (e.g. *EPO*) (Semenza, 2012). Besides tumor hypoxia or pVHL loss, other potential regulators of HIF1α may exist, which remains to be investigated.

In our current study, we investigated the role of ZHX2 as a new oncogene in TNBC, where it activates HIF1α signaling. In addition, we also provide some evidence for critical residues on ZHX2 that binds with DNA, which contributes to TNBC tumorigenicity.

## Results

### ZHX2 is Amplified in TNBC and is Potentially Regulated by pVHL

*ZHX2* is located on 8q24, where c-*Myc* resides. Analysis of copy number across different cancer types from The Cancer Genome Atlas (TCGA) showed that *ZHX2* is amplified in various cancers, including ovarian cancer (∼40%) and breast cancer (∼15%) (Figure 1A). Importantly, *ZHX2* and c-*Myc* share co-amplification in most cancer types observed (Figure 1B). Interestingly, *ZHX2* is not amplified in ccRCC (Referred as KIRC in Figure 1A, B), where it is regulated mainly post-transcriptionally by pVHL loss in this cancer (Zhang *et al*., 2018). Detailed analyses of several breast cancer patient datasets also revealed that TNBC had the highest ZHX2 amplification rate in all breast cancer subtypes (Figure 1C; Table Supplement 1). Next, we performed correlation studies to examine the copy number gain of *ZHX2* and its expression in TCGA (Cancer Genome Atlas, 2012) and METABRIC datasets (Curtis *et al*, 2012a) (Figure 1D). We found a significant correlation between copy number and gene expression (Figure 1D), suggesting that *ZHX2* amplification may be at least partially responsible for its overexpression in breast cancer. We further explored the effect of ZHX2 expression on breast cancer patient survival. ZHX2 overexpression (Affymetrix probe 236169_at) correlates with worse survival in TNBC but not in other breast cancer subtypes (Figure 1-figure supplement 1A). We obtained a panel of breast cancer cell lines as well as two immortalized normal breast epithelial cell lines, HMLE and MCF-10A. Interestingly, all breast cancer cell lines displayed relatively higher ZHX2 protein levels compared to HMLE or MCF-10A (Figure 1E). We also obtained 10 pairs of TNBC patient tumors and paired normal tissue. Consistently, ZHX2 was upregulated in the majority of tumors compared to normal (7 out of 10) (Figure 1F).

**Figure. 1.**
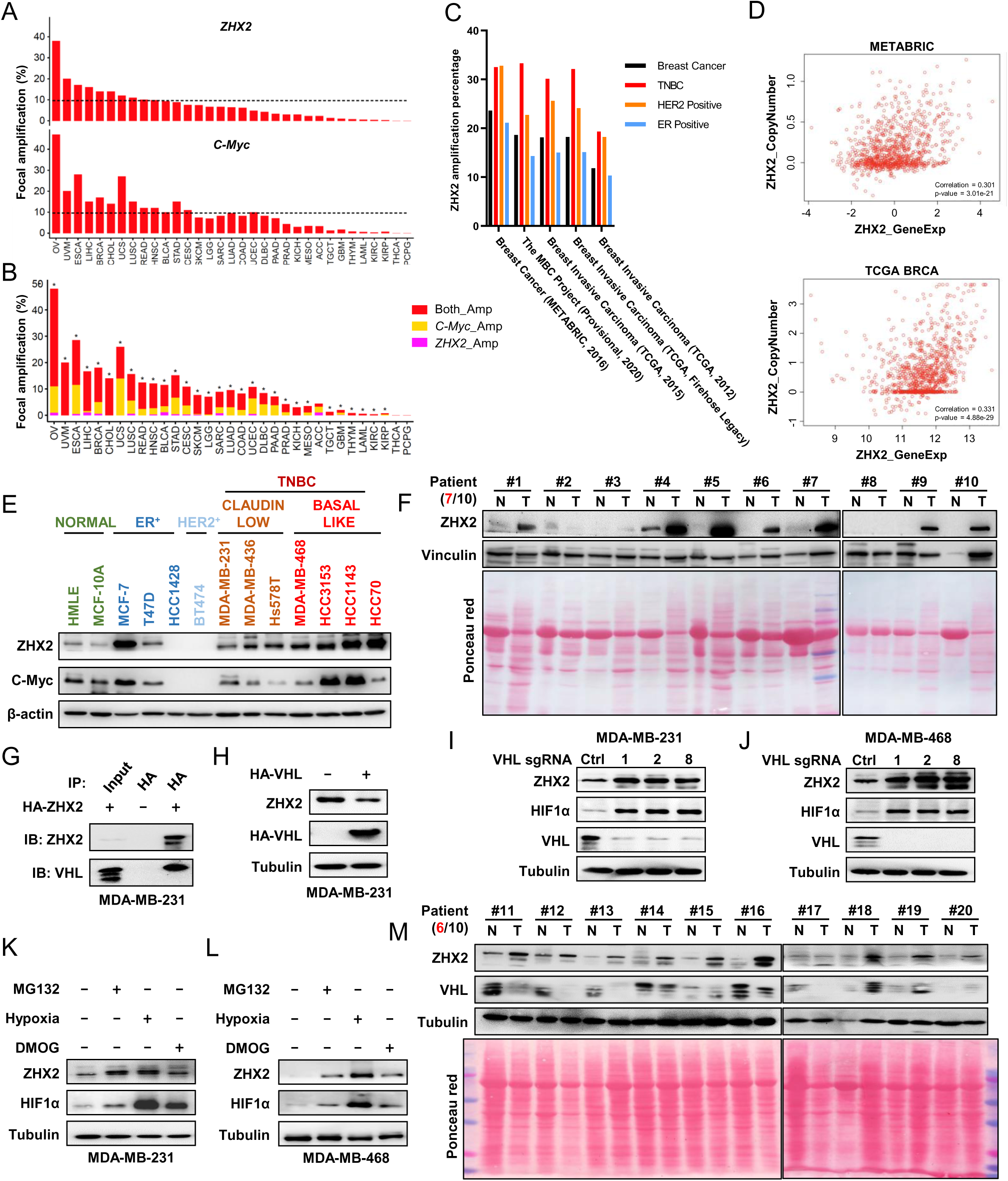
ZHX2 is amplified in TNBC and is potentially regulated by pVHL. (**A, B**) The percentage of tumor samples with ZHX2 (top) or MYC (bottom) focal amplification across cancer types (A) or samples with both ZHX2 and MYC (red), ZHX2- specific (magenta), and c-MYC-specific (orange) (B). Asterisk indicates statistical significance for overlap of ZHX2 and MYC focal amplification (Fisher’s exact test, p < 0.05). **(C)** The percentage of ZHX2 amplification of different breast cancer subtypes in several breast cancer datasets. **(D)** The relation of ZHX2 copy number gain and its expression in TCGA datasets and METABRIC datasets. **(E)** Immunoblots of lysates from different normal breast cell and breast cancer cell lines. **(F)** Immunoblots of lysates from paired TNBC patient-derived non-tumor (N) and tumor (T) breast tissues. **(G)** Immunoprecipitations (IP) of MDA-MB-231 cells infected with either control vector (EV) or FLAG-HA-ZHX2. **(H)** Immunoblots of lysates with indicated antibodies from MDA-MB-231 cells infected with either EV or HA-VHL. (**I, J**) Immunoblot of cell lysates from MDA-MB-231 (I) or MDA-MB-468 (J) infected with lentivirus encoding either VHL sgRNAs (1, 2, or 8) or control sgRNA (Ctrl). (**K, L**) Immunoblots of lysates from MDA-MB-231 (K) or MDA-MB-468 (L) cells treated with indicated inhibitors for 8 h. (**M**) Immunoblots of lysates from another 10 pair of TNBC patient-derived non-tumor (N) and tumor (T) breast tissues. **Figure supplement 1**. ZHX2 overexpression leads worse survival and is potentially regulated by pVHL in breast cancer. **Figure 1—source data.** Uncropped western blot images for Figure 1. **Figure supplement 1—source data.** Uncropped western blot images for Figure supplement 1.

Our previous research established an oncogenic role of ZHX2 in ccRCC as a pVHL substrate (Zhang *et al*., 2018). However, the role of ZHX2 in other cancers remains largely unclear. In addition, it is unclear whether ZHX2 may also act as a pVHL target in breast cancer. Since TNBC had the highest ZHX2 amplification rate in all breast cancer subtypes (Figure 1C; Table supplement 1), we decided to focus on TNBC for this current study. To this purpose, we first examined the relationship between the expression of ZHX2 and pVHL in several TNBC cell lines (MDA-MB-231, MDA-MB-436, MDA-MB-468, HCC3153 and HCC70) as well as two normal breast epithelial cell lines, HMLE and MCF-10A. Interestingly, all TNBC cell lines displayed relatively lower pVHL protein levels corresponding with higher ZHX2 protein level when compared to normal breast cells (Figure 1-figure supplement 1B). Our co-immunoprecipitation (Co-IP) experiments showed that ZHX2 interacts with pVHL in two representative TNBC cell lines (Figure 1G; Figure 1-figure supplement 1C). Next, we aimed to examine whether pVHL can promote the degradation of ZHX2, therefore decreasing ZHX2 protein levels in these cells. First, we overexpressed HA-VHL in two different TNBC cell lines and found that pVHL overexpression leads to decreased ZHX2 protein levels (Figure 1H; Figure 1-figure supplement 1D). Conversely, we also deleted pVHL expression by three independent sgRNAs (#1, 2, 8) in these two cell lines and found that pVHL depletion led to upregulation of ZHX2 protein (Figure 1I and J). Our previous research showed that ZHX2 regulation by pVHL potentially depends on ZHX2 prolyl hydroxylation (Zhang *et al*., 2018). We treated these two cell lines with hypoxia, the pan-prolyl hydroxylase inhibitor DMOG, or the proteasomal inhibitor MG132 and found that ZHX2 was upregulated by these inhibitors (Figure 1K and L), further strengthening the conclusion that ZHX2 is regulated by pVHL for protein stability through potential prolyl hydroxylation in breast cancer. In addition, we obtained 10 additional pairs of TNBC tumors and paired normal tissue to analyze ZHX2 and pVHL protein levels. In accordance with the cell line data, we found that ZHX2 was upregulated in most of tumor tissues compared to normal, coinciding with decreased pVHL protein levels in respective tumor tissues (Figure 1M). Our data suggest that ZHX2 may be regulated by pVHL and play an important role in TNBC.

### ZHX2 is Essential for TNBC Cell Proliferation and Invasion

Next, we examined the potential role of ZHX2 in TNBC cell proliferation and invasion. First, we obtained two previously validated ZHX2 shRNAs (sh43, sh45) (Zhang *et al*., 2018) and these shRNAs led to efficient ZHX2 protein (Figure 2A) and mRNA (Figure 2B) downregulation in both MDA-MB-231 and MDA-MB-468 cells. Next, we found that ZHX2 depletion led to decreased cell proliferation in both cell lines as a function of time, 2-D colony formation as well as 3-D soft agar growth (Figure 2C-F). One important contributor for the poor prognosis in TNBC is the aggressively invasive nature of this subtype. Therefore, we also used the Boyden chamber assay to examine the effect of ZHX2 on cell invasion in TNBC cells. Consistent with the results above, ZHX2 depletion led to decreased cell invasion in several different TNBC cell lines (Figure 2G and H). We also showed that ZHX2 shRNA induced phenotypes in TNBC cells could be completely rescued by shRNA-resistant ZHX2 (Figure 2I-N; Figure 2-figure supplement 2A-G), suggesting that these phenotypes were due to on-target depletion of ZHX2 by shRNAs.

**Figure. 2.**
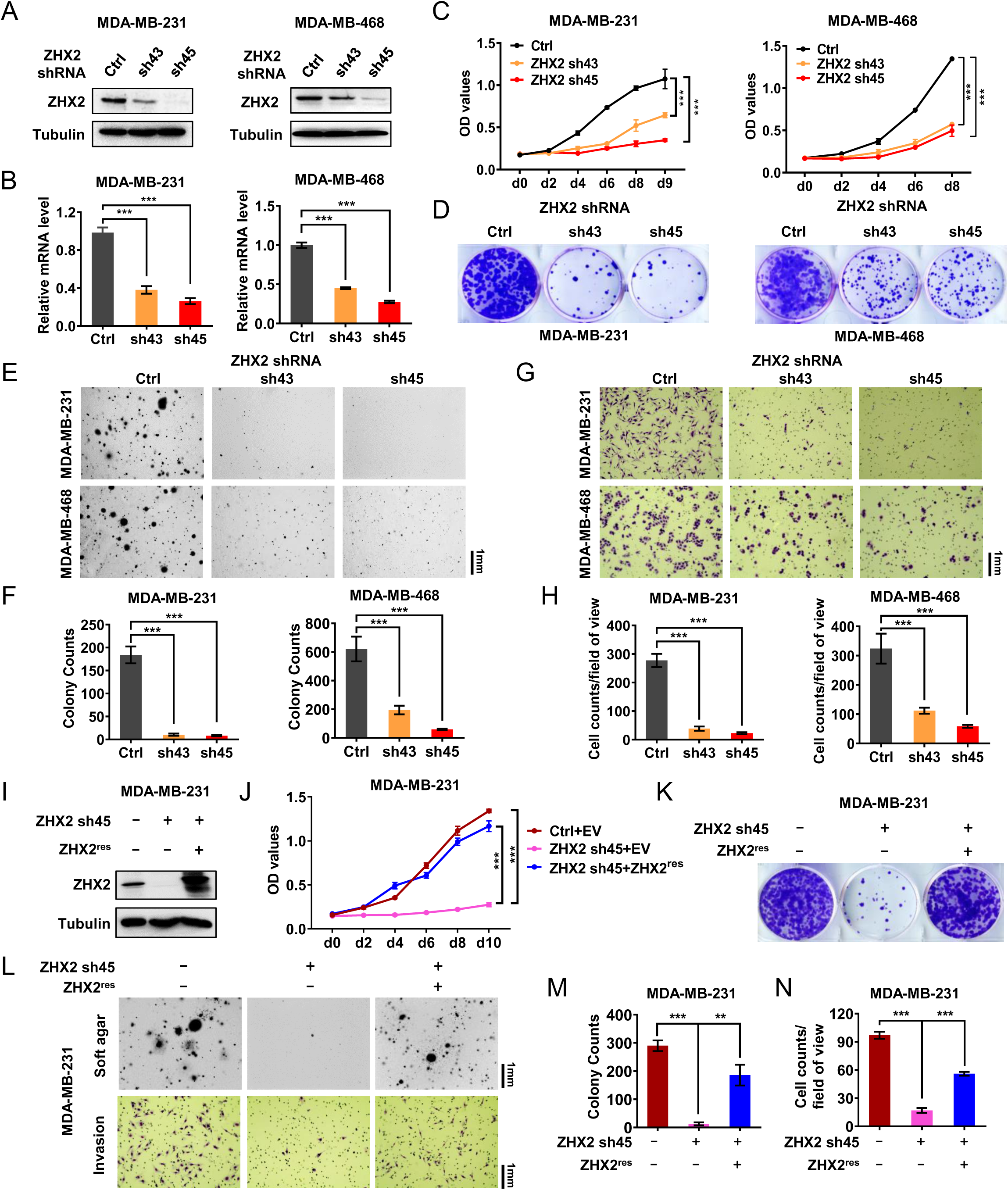
ZHX2 is essential for TNBC cell proliferation and invasion. (**A-H**) Immunoblot of cell lysates (A), qRT-PCR of RNA (B), cell proliferation assays (C), 2D-clones (D), soft agar growth (E) and quatification (F), invasion (G) and quantification (H) of MDA-MB-231/468 cells infected with lentivirus encoding two individual ZHX2 shRNAs (43, 45) or control shRNA (Ctrl). (**I-N**) Immunoblot of cell lysates (I), cell proliferation (J), 2D-clones (K), soft agar growth (upper) and invasion assays (down) (L) as well as quantification (M-N) of MDA-MB-231 cells infected with lentivirus encoding ZHX2 sh45-resistant FLAG-HA-ZHX2 or control vector (EV), followed by ZHX2 sh45 or Ctrl. Error bars represent mean ± SEM, unpaired t-test. * denotes p value of <0.05, * * denotes p value of <0.01, * * * denotes p value of <0.005. **Figure supplement 2**. The phenotype of ZHX2 shRNA on cell proliferation and invasion is due to its ontarget effect. **Figure 2—source data.** Uncropped western blot images for Figure 2. **Figure supplement 2—source data.** Uncropped western blot images for Figure supplement 2.

### ZHX2 is Important for TNBC Cell Proliferation *in Vivo*

Next, to examine the ability of ZHX2 to maintain TNBC tumor growth *in vitro* and *in vivo*, we first infected two TNBC cell lines with doxycycline inducible ZHX2 shRNAs (Teton sh43, sh45). Upon doxycycline addition, we achieved efficient depletion of ZHX2 protein levels, corresponding to decreased ZHX2 mRNA levels (Figure 3A; Figure 3-figure supplement 3A and B) in the cells. Next, MTS assays showed that ZHX2 depletion upon doxycycline addition led to decreased TNBC cell proliferation, 2-D growth, 3-D anchorage independent growth (Figure 3B-E; Figure 3-figure supplement 3C-F). Next, we expressed firefly luciferase in the doxycycline-inducible ZHX2 shRNA (sh43 or sh45) or control shRNA cells before injecting these cells orthotopically into the 4^th^ mammary fat pads of NoD SCID Gamma (NSG)-deficient mice. We performed weekly bioluminescence imaging to ensure the successful implantation and growth of TNBC tumor cells. After 12 days post-implantation when palpable tumors were formed, we fed these mice doxycycline chow. We found that ZHX2 depletion by both hairpins significantly decreased tumor growth overtime. Upon necropsy, ZHX2 shRNA-infected TNBC cells displayed reduced tumor burden retrieved from tumor-bearing mice compared to control mice (Figure 3F and G). We also performed a western blot for tumors extracted from the mammary fat pad and found decreased ZHX2 protein levels in ZHX2 shRNA-infected groups (Figure 3H), arguing that anti-tumor effect in these groups may result from efficient ZHX2 knockdown in these TNBC cells. In addition, we also measured spontaneous lung metastasis *ex vivo* upon necropsy and found that ZHX2 depletion led to significantly decreased lung metastasis in TNBC (Figure 3I and J). Lastly, we also injected ZHX2 Teton sh45-infected cells into the mammary fat pad, followed by regular chow or doxycycline chow. Consistent with results above, doxycycline chow led to significantly decreased TNBC tumor growth over time (Figure 3K and L) corresponding with lower ZHX2 protein levels in these tumors (Figure 3M). Taken together, our data strongly indicate that ZHX2 is important for TNBC tumorigenesis *in vivo*.

**Figure. 3.**
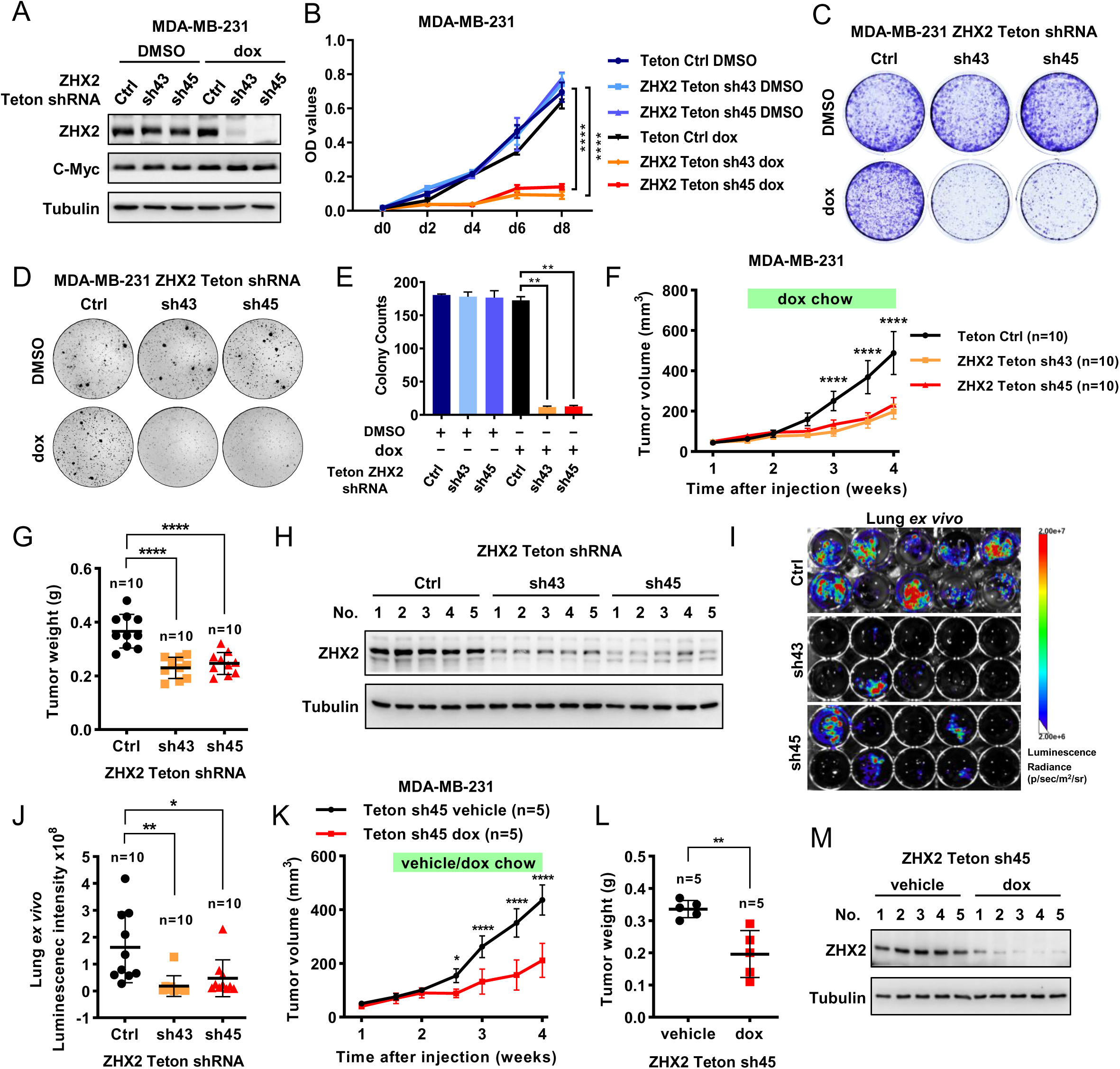
ZHX2 is important for TNBC cell proliferation *in vivo*. (**A-E**) Immunoblot of cell lysates (A), cell proliferation (B), 2D-colony growth (C), soft agar growth (D) and quantification (E) of MDA-MB-231 cells infected with lentivirus encoding Teton-ZHX2 shRNAs (43, 45) or Teton-Ctrl. (**F-G**) Tumor growth (F), and tumor weight (G) of doxycycline-inducible ZHX2 knockdown MDA-MB-231 cells injected orthotopically at the mammary fat pad of NSG mice. Treatment of doxycycline food started as indicated time. (**H**) Immunoblot of tumor lysates from mouse that injected doxycycline-inducible ZHX2 knockdown MDA-MB-231 cells. (**I, J**) Images (I) and plot (J) of lung necropsy of doxycycline-inducible ZHX2 knockdown MDA-MB-231 cells injected orthotopically at the mammary fat pad of NSG mice. (**K, L**) Tumor growth (K), and tumor weight (L) of doxycycline induced or not induced ZHX2 sh45 MDA-MB-231 cells injected orthotopically at the mammary fat pad of NSG mice. (**M**) Immunoblot of tumor lysates from mouse treated with regular chow or doxycycline chow. Error bars represent mean ± SEM, unpaired t-test. * denotes p value of <0.05, * * denotes p value of <0.01, * * * denotes p value of <0.005. **Figure supplement 3**. ZHX2 is important for maintaining TNBC tumorigenesis in vivo. **Figure 3—source data.** Uncropped western blot images for Figure 3. **Figure supplement 3—source data.** Uncropped western blot images for Figure supplement 3.

### ZHX2 Regulates HIF Signaling in TNBC

Next, we aimed to determine the molecular mechanism by which ZHX2 contributes to TNBC. For this purpose, we performed RNA-seq analyses in MDA-MB-231 cells with two independent ZHX2 shRNAs (sh43, sh45) and found that two individual shRNAs concordantly altered gene expression patterns (Figure 4A). Pathway enrichment as well as gene set enrichment analyses (GSEA) revealed that genes differentially expressed following ZHX2 depletion were enriched for members of the hypoxia pathway (Figure. 4B; Figure 4-figure supplement 4A-C). Overall, there were 7,690 genes differentially expressed following ZHX2 silencing by shRNA, and 1,849 (24%) of these genes overlapped with genes differentially expressed in a previously published HIF double knockout (HIF1α and HIF2α double knockout, HIF DKO) model (Figure 4C) (Chen *et al*, 2018). Since ZHX2 was similarly reported as an oncogene in ccRCC and preferentially upregulated the transcription of downstream genes (Zhang *et al*., 2018), we focused on the 3,969 ZHX2 positively regulated genes (downregulated following ZHX2 silencing by shRNA) as these genes may be more relevant in breast cancer. Of these, 678 genes (17%) overlapped with downregulated genes in the HIF DKO RNA-seq (Figure 4B-D), representing a significant association (adj. p = 1.07 x 10^-69^). This data strengthens the potential functional link between HIF and ZHX2. The genes positively regulated by ZHX2 were also enriched for other potentially relevant biological pathways such as cell adhesion and cell morphogenesis (Figure 4B). Next, we examined whether ZHX2 depletion affected some of the canonical HIF target genes. Based on our ZHX2 RNA-seq and GSEA results (Figure 4-figure supplement 4C), we chose a few representative HIF target genes and examined their expression levels by qRT-PCR in MDA-MB-231 cells infected with control or ZHX2 shRNA under either normoxia or hypoxia. As expected, hypoxia treatment increased the expression of HIF target genes, and this effect was ameliorated by ZHX2 depletion (Figure 4-figure supplement 4D), suggesting that ZHX2 affects HIF activity and regulates HIF target gene expression in TNBC. Consistently, HIF reporter assay found that ZHX2 depletion led to decreased HIF reporter activity either under normoxia or hypoxia (Figure 4-figure supplement 4E).

**Figure. 4.**
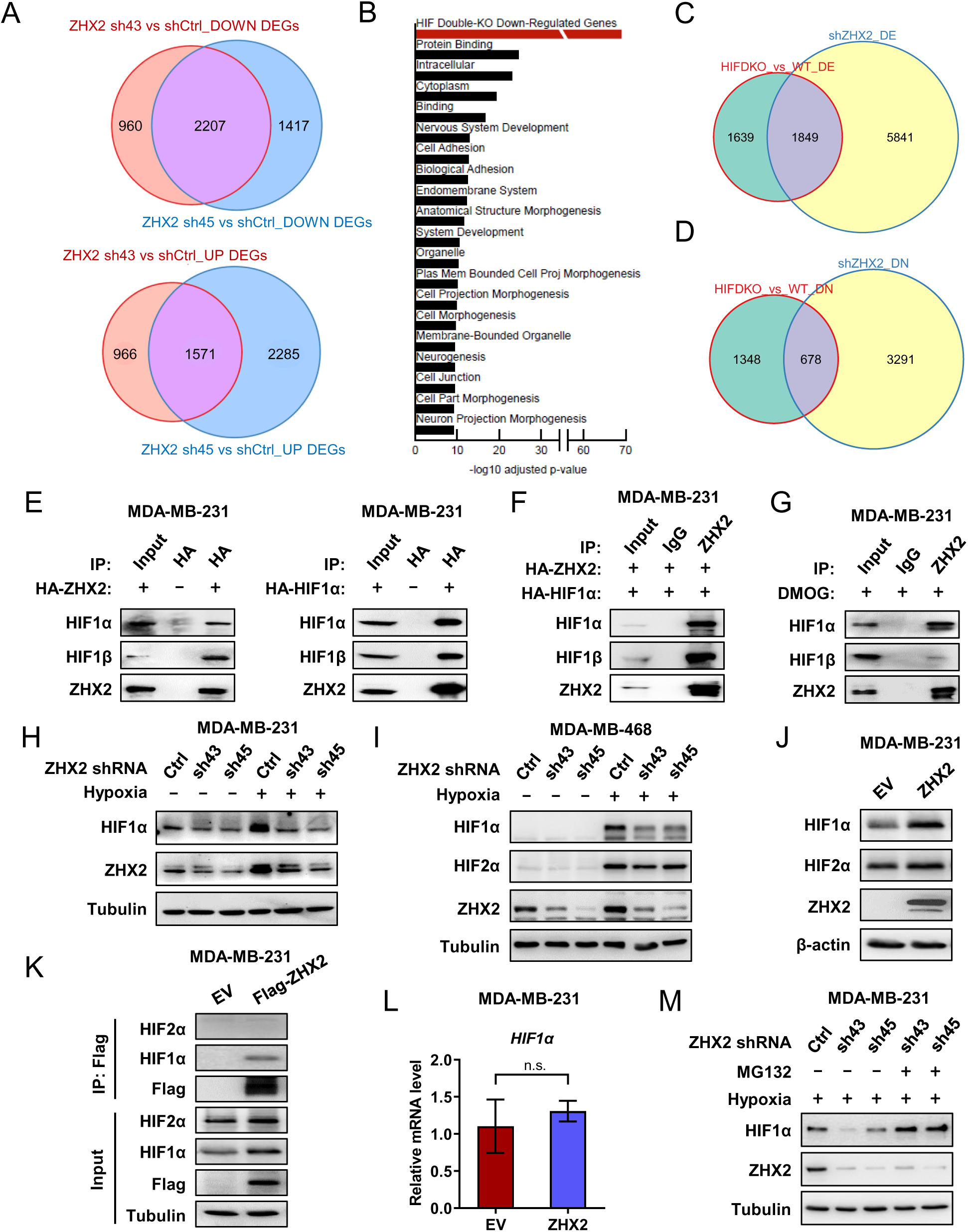
ZHX2 regulates HIF signaling in TNBC. **(A)** Venn diagram showing overlap in downregulated (top) and upregulated (bottom) genes between two different ZHX2 shRNA. **(B)** Pathway analysis of the significantly decreased pathways in ZHX2 depleted MDA-MB-231 cells. (**C, D**) Venn diagram showing overlap in differentially expressed genes (C) and downregulated genes (D) between ZHX2 depletion and HIF double knock-out (DKO) (GSE108833). (**E, F**) Immunoblots of immunoprecipitations (IP) of MDA-MB-231 cells overexpress either FLAG-HA-ZHX2 or HA-HIF1α (E) or both ZHX2 and HIF1α (F). (**G**) Immunoblots of immunoprecipitations (IP) of MDA-MB-231 cells treated with DMOG for 8 hours. (**H, I**) Immunoblots of cell lysates from MDA-MB-231 cells (H) and MDA-MB-468 cells (I) infected with lentivirus encoding ZHX2 shRNAs or Ctrl, followed treated with normoxia or hypoxia (1% O_2_). (**J, L**) Immunoblots (J) and immunoprecipitations (K) of cell lysates, qRT-PCR of mRNA (L) from MDA-MB-231 cells infected with lentivirus encoding EV or FLAG-HA-ZHX2. (**M**) Immunoblots of cell lysates from MDA-MB-231 cells infected with lentivirus encoding ZHX2 shRNAs or Ctrl treated with MG132 overnight under hypoxia (1% O_2_). Error bars represent mean ± SEM, unpaired *t*-test. * denotes p value of <0.05, * * denotes p value of <0.01, * * * denotes p value of <0.005. **Figure supplement 4**. ZHX2 regulates HIF1 signaling in TNBC. **Figure 4—source data.** Uncropped western blot images for Figure 4. **Figure supplement 4—source data.** Uncropped western blot images for Figure supplement 4.

Next, to gain further insight on how ZHX2 affect the HIF signaling. Consider a well characterization on HIF1 function in TNBC tumorigenesis by previous studies (Bos *et al*, 2001; Briggs *et al*., 2016). We first performed Co-IP experiments and showed that ZHX2 bound with HIF1α and HIF1β exogenously as well as endogenously (Figure 4E-G). In addition, ZHX2 knockdown led to decreased HIF1α protein levels in two TNBC cell lines under hypoxia condition (Figure 4H and I). Conversely, ZHX2 overexpression led to increased HIF1α protein levels under normoxia (Figure 4J). However, these ZHX2 loss-of-function or gain-of-function manipulations did not grossly change HIF2α protein level (Figure 4I and J). Co-IP experiments showed that ZHX2 could not bind with HIF2α (Figure 4K). These results suggest that ZHX2 regulates the hypoxia signaling mainly through HIF1α. Interestingly, qRT-PCR showed that ZHX2 overexpression did not increase HIF1α mRNA level (Figure 4L), suggesting a mechanism of post-transcriptional regulation on HIF1α We then treated the ZHX2 knockdown cells with the proteasome inhibitor MG132 under hypoxia, which could fully rescue the HIF1α protein level in the knockdown cells (Figure 4M). Altogether, these data suggested ZHX2 controls HIF1α protein stability by preventing proteasome-mediated degradation. On the other hand, western blot analysis showed that HIF1α knockdown did not affect ZHX2 protein levels (Figure 4-figure supplement 4F). It is important to note that the detailed mechanism on how ZHX2 binds with HIF1 and transactivates HIF1 signaling remains unclear, which awaits future investigation.

To identify the direct downstream target genes of ZHX2 and HIF that may be important in TNBC, we performed chromatin immunoprecipitation followed by high-throughput sequencing (ChIP-seq) to assess the genomic binding pattern of ZHX2 in TNBC. We identified 957 binding sites across the genome, of which 94% of them overlap with H3K27ac and 96% of them overlap with H3K4me3 (Figure 5A), indicating these overlapping peaks bound preferentially to active promoters for gene expression (Shlyueva *et al*, 2014). We again focused on the ZHX2 positively regulated genes (downregulated following ZHX2 silencing by shRNA) that exhibited ZHX2 binding in the promoter (transcription start site ± 5 kb). These promoters (n = 258) demonstrated robust enrichment for HIF1α (Chen *et al*., 2018), as well as H3K4me3 and H3K27ac (Rhie *et al*, 2014) (Figure 5A). We then filtered these genes further, focusing on only those bound and positively regulated by HIF1α and identified seven interesting candidate genes for functional validation and follow-up (*PTGES3L*, *KDM3A*, *WSB1*, *AP2B1*, *OXSR1*, *RUNDC1* and *COX20*). We performed qRT-PCR analysis and found that ZHX2 depletion by shRNAs indeed led to decreased mRNA expression for all of these target genes (Figure 5B). Conversely, we also overexpressed ZHX2 in TNBC cells and found that ZHX2 overexpression led to increased expression of these target genes (Figure 5C), arguing that ZHX2 promotes HIF signaling by at least directly activating these targets in TNBC. To further strengthen whether these are HIF downstream target genes, we also obtained HIF DKO cells and found that HIF depletion led to decreased expression of these target genes under hypoxic conditions (Figure 5D). Taken together, our data suggests that ZHX2 binds with HIF and affects HIF activity in TNBC.

**Figure. 5.**
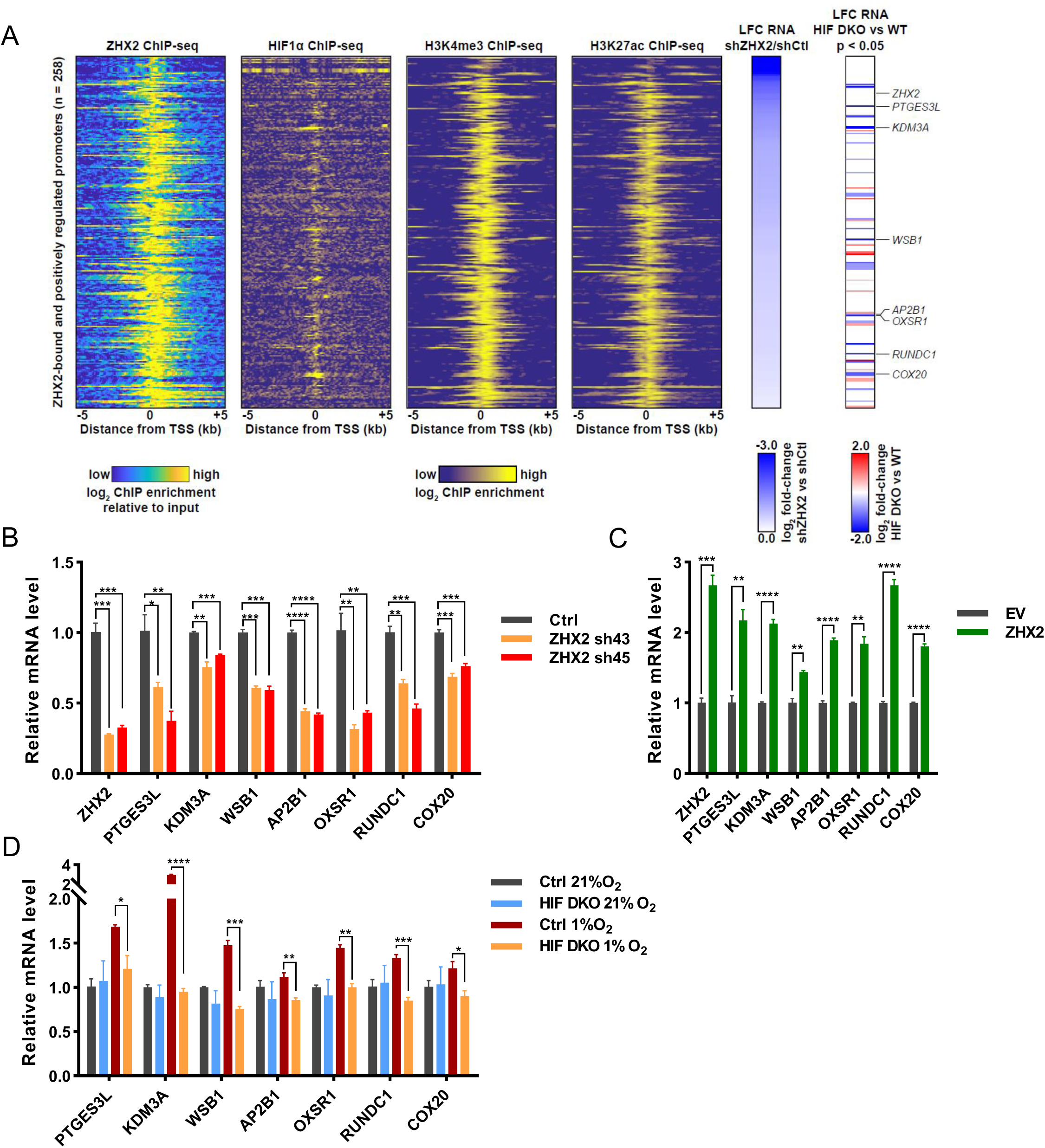
Representative ZHX2 and HIF downstream targets and analysis of their chromatin binding motifs. (**A**) Integrated analyses of ChIP-Seqs (including ZHX2, HIF1α, H3K4me3 and H3K27ac), signals expressed as relative to input control when available. Log2 fold change (LFC) for ZHX2 knock down RNA-Seq and HIF double knockout (HIF DKO) RNA-Seq; Critical target genes were marked on the right. (**B-D**) qRT-PCR quantification of ZHX2 target genes from MDA-MB-231 cells infected with lentivirus encoding ZHX2 shRNAs (43, 45) (B), EV or FLAG-HA-ZHX2 (C) or HIF double knockout under normoxia (21% O_2_) and hypoxia (1% O_2_) (D).

### Potential Important Sites on ZHX2 that May affect its Function in TNBC

Previous research showed that ZHX2 contributed to ccRCC tumorigenesis by at least partially activating NF-κB signaling (Zhang *et al*., 2018). In that setting, ZHX2 may act as transcriptional activator by overlapping primarily with H3K4me3 and H3K27Ac epigenetic marks. However, it remains unclear whether there may be critical residues on ZHX2 that may mediate its binding to DNA and exert its transcriptional activity. To this end, we conducted both data-driven analyses and structural simulations to predict the essential DNA binding residues. Detailed methods are described in Supplementary. Briefly, many DNA-bound structures of human HD2, 3 and 4 (PDB ID: 3NAU, 2DMP and 3NAR) were homology-modeled by SWISS-MODEL (23) using as many X-ray/NMR-solved homologous DNA-bound HD proteins as the structural templates. 12, 20 and 15 DNA-bound complexes were modeled for HD2, HD3 and HD4, respectively (Figure 5-figure supplement 5). We then counted the number of DNA-protein contacts at the atomic level for each residue in the DNA-contacting helices in HDs, normalized by the number of DNA-complexed structures used for each HD. The top-ranked DNA-contacting residues in HDs are listed together with their evolutionary conservation in Table supplement 2. We found that Lys485/Arg491 in HD2, Arg581 in HD3 and Arg674 in HD4 are the most contacted residues in DNA binding, where Arg674 receives the highest contact among all the HD proteins. Our MD simulations further revealed that Arg491 in HD2, Arg581 in HD3 and Arg674 in HD4 indeed have the highest affinity with DNA, in terms of MM/PBSA-derived contact potential energies (see Table supplement 3 and 4; Movie Supplement1, 2 and 3), among other residues in the same proteins. Among the top 4 DNA-contacting residues in each of the HD proteins, we consider those with relatively high sequence conservation being important for ZHX2 binding to DNA, which may affect the phenotype of TNBC. These residues are Asp489 (D489), Arg491 (R491), Glu579 (E579), Arg581 (R581), Lys582 (K582), Arg674 (R674), Glu678 (E678), and Arg680 (R680) (Table Supplement 2).

Given this, we generated a series of TNBC breast cancer cell lines where we depleted endogenous ZHX2 expression and restored with exogenous shRNA-resistant ZHX2 WT or mutant versions (D489A, R491A, E579A, R581A, K582, R674A, E678A, or R680A). First, upon generation of stable cell lines, we performed western blot analyses and confirmed that these cell lines all expressed similar amounts of ZHX2, relatively comparable to endogenous ZHX2 levels in MDA-MB-231 cells (Figure 6A). Next, we performed 2-D cell proliferation MTS assays. Our cell proliferation data showed that consistent with our previous results, ZHX2 depletion led to decreased TNBC cell proliferation, and this phenotype was completely rescued by WT ZHX2. On the other hand, some of mutants (including R581A and R674A) failed to rescue the cell growth defect in ZHX2 shRNA infected MDA-MB-231 cells (Figure 6B).

**Figure. 6.**
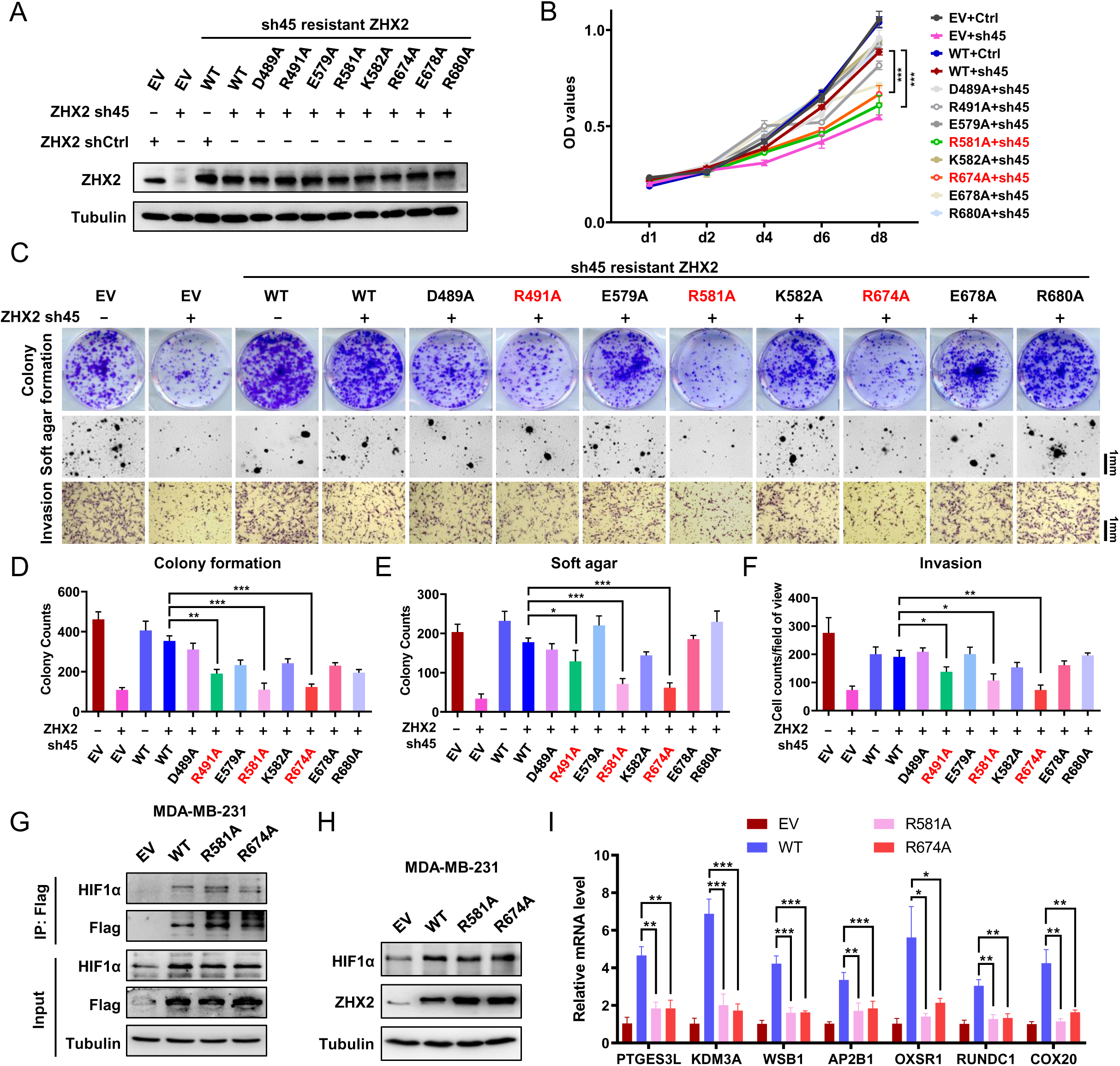
Identification of important sites on ZHX2 that may affect its function in TNBC. (**A-F**) Immunoblot of cell lysates (A), cell proliferation (B), 2D colony (top), 3D soft-agar (middle) and invasion (bottom) (C) as well as quantification 2D colony (D), 3D soft-agar (E) and invasion (F) of MDA-MB-231 cell lines infected with lentivirus encoding either ZHX2 wild type (WT) or mutation, followed by ZHX2 sh45 or Ctrl. (**G-I**) Immunoprecipitations (G), Immunoblot (H) of cell lysates and qRT-PCR of mRNA (I) of MDA-MB-231 cells infected with lentivirus encoding either ZHX2 WT or mutation. Error bars represent mean ± SEM, n=3 replicates per group, unpaired *t*-test. *** denotes p value of <0.005. **Figure supplement 5**. Residue based DNA contact analysis derived from ensembles of DNA bound HD proteins. **Figure 6—source data.** Uncropped western blot images for Figure 6.

Motivated by our cell proliferation assay results, we further examined cell proliferation phenotypes using long-term 2-D colony formation, 3-D anchorage independent growth and cell invasion assays. ZHX2 R491A, R581A and R674A mutants displayed a defect in cell growth in TNBC cell lines (Figure 6C-F). In summary, our results showed that there may be multiple residues (including R491, R581 and R674) that may be important in regulating the phenotype of ZHX2 in TNBC. Co-IP analysis showed that mutated ZHX2 (R581A and R674A) can interact with HIF1α as wild type ZHX2 (Figure 6G). While western blot analysis showed that the ZHX2 mutations did not affect the HIF1α protein levels (Figure 6H), qRT-PCR showed that the ZHX2 mutations indeed led to decreased mRNA expression of HIF1α targeted genes (Figure 6I). Taken together, our data suggests that these residues are essential for mediating the transactivation of ZHX2 on HIF1α target genes.

## Discussion

In this study, we discover that ZHX2 is an important oncogene in TNBC. Depletion of ZHX2 leads to decreased TNBC cell proliferation as well as invasion. By performing gene expression analyses, ZHX2-regulated genes display a significant overlap with HIF-regulated genes in TNBC. ZHX2 binds with HIF1α and HIF1β and regulates HIF1α protein levels and transcriptional activity. By using structural simulation and the re-constitution system, we pinpoint residues (R491, R581 and R674) on ZHX2 that may be important for its DNA binding function as well as tumorigenic potential. Overall, our study establishes an important role of ZHX2 in regulating HIF1α signaling and tumorigenesis in TNBC.

From a genomic perspective, it is known that *ZHX2* is located on 8q24, a chromosomal region frequently amplified in cancers. Indeed, *ZHX2* is amplified in several cancers, including breast cancer, ovarian cancer, and prostate cancer. In most cases, ZHX2 is co-amplified with another well-established oncogene c-*Myc* (Figure 1A-C). This finding bears several implications. First, it suggests that ZHX2 and c-Myc may act in concert in promoting tumorigenesis. Second, the role of ZHX2 in cancers can be context dependent. Although ZHX2 may be amplified in multiple cancers, its protein levels can be regulated post-transcriptionally. Our previous research showed that in ccRCC, ZHX2 can be regulated by pVHL potentially through hydroxylation on multiple proline residues in the PRR domain (Zhang *et al*., 2018). Therefore, the presence or absence of factors that mediate prolyl hydroxylation in the same niche as ZHX2 can dictate its regulation and downstream function. Further, it remains uncertain whether ZHX2 interacts with DNA directly or indirectly via other transcription factors to exert its transcriptional regulation on downstream target genes. The repertoire of different co-activators/repressors with which ZHX2 may interact can therefore govern its localization in the genome and thus its downstream function in different cancer settings.

Thus far, it remains unclear which residues on ZHX2 are critical for its transcriptional activity as well as its oncogenic role in cancer. We also did MD simulation to confirm the import residues for regulating the phenotype of ZHX2. We found that the top DNA-contacting residues across the three HDs were arginine namely Arg491, Arg581, and Arg674. In terms of interaction, it can be observed that Arg491 and Arg581 of HD 2 and 3, respectively, were seen to be interacting with the DNA’s phosphate backbone throughout the 350ns MD simulations (Movie supplement 1-2). However, in the case of HD 4, Arg674 was seen to be interacting first with the nucleobases within 4Å distance. Shortly after ∼120ns, the DNA slowly drifted away but this was prevented because of an interaction between Arg674 and the DNA’s phosphate backbone (Movie supplement 3). These arginine-DNA phosphate backbone interactions could be critical for the stabilization of transcription factor binding with the DNA to either stimulate or repress the transcription of a specific gene. By performing ZHX2 depletion and reconstitution experiments, we found that several residues located in the HDs of ZHX2 may be important for its oncogenic role in TNBC (Figure 6A-F). In addition, these three arginine residues are found to be important on mediating the effect of ZHX2 on transactivating HIF1α activity in TNBC cells (Figure 6I). Further research needs to be performed to determine whether these residues are critical for ZHX2 localization to DNA, either directly or indirectly via recruitment of co-activators/repressors. Lastly, given that we already found three residues important for the function of ZHX2 in TNBC, we can potentially design small peptides to competitively bind ZHX2 and inhibit its localization to DNA. By engineering these peptides to be membrane permeable, we can potentially test whether they can inhibit the oncogenic role of ZHX2 in TNBC. Given that ZHX2 inhibitors are still not available at this time, these peptide inhibitors can be used as a proof-of-principle approach to motivate further development of specific ZHX2 inhibitors in potential cancer therapies.

## Materials and Methods

### Cell Culture and Reagents

MDA-MB-231, MDA-MB-436, MCF-7, Hs578T and 293T cells were cultured in Dulbecco’s Modified Eagle’s Medium (DMEM) (GIBCO 11965118) supplemented with 10% fetal bovine serum (FBS) and 1% penicillin-streptomycin (Pen Strep). T47D, BT474, HCC1428, HCC3153, HCC1143, HCC70 and MDA-MB-468 cells were cultured in 10% FBS, 1% Pen Strep RPMI 1640 (GIBCO 11875093). Normal breast epithelial cells HMLE and MCF-10A were cultured in MEGM (Lonza CC-3151) containing SingleQuots Supplements (Lonza CC-4136). 293T cells were obtained from UNC Tissue Culture Facility and authenticated by short tandem repeat testing. HMLE and HCC3153 are in-house cell lines. All other cell lines were obtained from ATCC. Mycoplasma detection was routinely performed to ensure cells were not infected with mycoplasma by using MycoAlert Detection kit (Lonza, LT07-218). Cells were maintained at 37°C in a 5% CO_2_ incubator. Cells were incubated overnight in 1% O_2_ hypoxia chamber for hypoxia treatment. Doxycycline (D9891) was purchased from Sigma-Aldrich, DMOG (D1070-1g) was from Frontier Scientific, and MG132 (IZL-3175-v) was from Peptide International.

### Amplification status of ZHX2 in different breast cancer subtype

All data in table S4 were got from cBioPortal (https://www.cbioportal.org/) (Gao *et al*, 2013). We searched the percentage of ZHX2 amplification in all breast cancer datasets, and found ZHX2 was mainly amplified in seven datasets: Breast Cancer (METABRIC, Nature 2012(Curtis *et al*, 2012b) & Nat Commun 2016(Pereira *et al*, 2016)), The Metastatic Breast Cancer Project (Provisional, February 2020), Breast Invasive Carcinoma (TCGA, Cell 2015)(Ciriello *et al*, 2015), Breast Invasive Carcinoma (TCGA, Firehose Legacy), Breast Invasive Carcinoma (TCGA, PanCancer Atlas), Metastatic Breast Cancer (INSERM, PLoS Med 2016)(Lefebvre *et al*, 2016), and Breast Invasive Carcinoma (TCGA, Nature 2012)(CancerGenomeAtlasNetwork, 2012). Two datasets, Breast Invasive Carcinoma (TCGA, PanCancer Atlas) and Metastatic Breast Cancer (INSERM, PLoS Med 2016)(Lefebvre *et al*., 2016), did not show the ER, PR and HER2 status, and were excluded from our study. In all the datasets, ER and PR status were determined by immunohistochemistry (IHC). Three datasets, Breast Cancer (METABRIC, Nature 2012(Curtis *et al*., 2012b) & Nat Commun 2016(Pereira *et al*., 2016)), The Metastatic Breast Cancer Project (Provisional, February 2020), and Breast Invasive Carcinoma (TCGA, Cell 2015)(Ciriello *et al*., 2015) assigned the HER2 status by the original researchers. Two datasets, Breast Invasive Carcinoma (TCGA, Cell 2015)(Ciriello *et al*., 2015) and Breast Invasive Carcinoma (TCGA, Firehose Legacy) assigned the HER2 status by two standard, IHC and fluorescence in situ hybridization (FISH). In this study, HER2+ status in these two datasets, were determined by IHC.

### Immunoblotting and Immunoprecipitation Experiments

EBC buffer (50mM Tris-HCl pH8.0, 120 mM NaCl, 0.5% NP40, 0.1 mM EDTA and 10% glycerol) supplemented with complete protease inhibitor and phosphoSTOP tablets (Roche Applied Bioscience) was used to harvest whole cell lysates at 4°C. Cell lysate concentrations were measured by Protein assay dye (Bio Rad). An equal amount of cell lysates was resolved by SDS-PAGE. For immunoprecipitation, whole-cell lysates were prepared in EBC buffer supplemented with protease inhibitor and phosphatase inhibitor. The lysates were clarified by centrifugation and then incubated with primary antibodies or HA antibody conjugated beads (HA beads, Roche Applied Bioscience) overnight at 4°C. For primary antibody incubation, cell lysates were incubated further with protein G sepharose beads (Roche Applied Bioscience) for 2 hours at 4°C. The bound complexes were washed with EBC buffer 5× times and were eluted by boiling in SDS loading buffer. Bound proteins were resolved in SDS-PAGE followed by immunoblotting analysis.

### Antibodies

Antibodies used for immunoblotting, immunoprecipitation and IHC staining were as follows: Rabbit anti ZHX2 antibody (Genetex, 112232), Rabbit anti HIF1α (Cell Signaling, 36169), Rabbit anti HIF1β (Cell Signaling, 5537), Rabbit anti VHL (Cell Signaling, 68547), rabbit anti HA tag (Cell Signaling, 3724), mouse anti α-Tubulin (Cell Signaling, 3873). Peroxidase conjugated goat anti-mouse secondary antibody (31430) and peroxidase conjugated goat anti-rabbit secondary antibody (31460) were from Thermo Scientific.

### Plasmids

pBABE HA-VHL, pcDNA-3.1-FLAG-HA-ZHX2(WT), pcDNA-3.1-FLAG-HA-ZHX2(ZHX2sh45 resistant), and pcDNA-3.1-HA-HIF1α were previously described. pcDNA-3.1-FLAG-HA-ZHX2(D489A), pcDNA-3.1-FLAG-HA-ZHX2(R491A), pcDNA-3.1-FLAG-HA-ZHX2(E579A), pcDNA-3.1-FLAG-HA-ZHX2 (R581A), pcDNA-3.1-FLAG-HA-ZHX2 (K582A), pcDNA-3.1-FLAG-HA-ZHX2 (R674A), pcDNA-3.1-FLAG-HA-ZHX2 (E678A), and pcDNA-3.1-FLAG-HA-ZHX2 (R680A) were constructed using standard molecular biology techniques. Quick Change XL Site-Directed Mutagenesis Kit (200516, Agilent Technologies) was used to construct ZHX2 mutants. The GATEWAY Cloning Technology (11789020 and 11791019, Invitrogen) was used to recombine plasmids for virus production. All plasmids were sequenced to confirm validity.

### Lentiviral shRNA, and sgRNA Vectors

Lentiviral ZHX2 shRNAs (pLKO vector based) were obtained from Broad Institute TRC shRNA library. sgRNAs were cloned into the lentiCRISPR v2 backbone (Addegene Plasmid #52961). Target sequences were as follows:

Control shRNA: AACAGTCGCGTTTGCGACTGG
ZHX2 shRNA (43): CCCACTAAATACTACCAAATA
ZHX2 shRNA (45): CCGTAGCAAGGAAAGCAACAA
HIF1α shRNA (3809): CCAGTTATGATTGTGAAGTTA
HIF1α shRNA (3810): GTGATGAAAGAATTACCGAAT
Control sgRNA: GCGAGGTATTCGGCTCCGCG
VHL sgRNA (1): CATACGGGCAGCACGACGCG
VHL sgRNA (2): GCGATTGCAGAAGATGACCT
VHL sgRNA (8): ACCGAGCGCAGCACGGGCCG

### Virus Production and Infection

293T packaging cell lines were used for lentiviral amplification. Lentiviral infection was carried out as previously described (Zhang *et al*., 2018). Briefly, viruses were collected at 48 h and 72 h post-transfection. After passing through 0.45-µm filters, viruses were used to infect target cells in the presence of 8 µg/mL polybrene. Subsequently, target cell lines underwent appropriate antibiotic selection.

### Cell Viability Assay

For MTS assay, cells were seeded in triplicate in 96-well plates (1000 cells/well) in appropriate growth medium. At indicated time points, cells were replaced with 90 µl fresh growth medium supplemented with 10 µl MTS reagents (Abcam, ab197010), followed by incubation at 37°C for 1-4 hrs. OD absorbance values were measured at 490 nm using a 96-well plate reader (BioTek).

### 2-D Cell Proliferation Assay

For colony formation assays, cells were seeded in duplicate in 6-well plates (2 x 10^3^ cells/well) in appropriate growth medium. Media was changed every two days. After 7 days, cells were fixed with 4% formaldehyde for 10 minutes at room temperature, stained for 10 minutes with 0.5% crystal violet and then washed several times with distilled water. Once dried, the plates were scanned.

### 3-D Anchorage Independent Soft Agar Growth Assay

Cells were plated in a top layer at a density of 10,000 cells per ml in complete medium with 0.4% agarose (Life Technologies, BP165-25), onto bottom layers composed of medium with 1% agarose followed by incubation at 4°C for 10 minutes. Afterwards, cells were moved to a 37°C incubator. Every 4 days, 200μl of complete media were added onto the plate. After 2-4 weeks, the extra liquid on the plate was aspirated, and 1 ml medium supplemented with 100 μg/ml iodonitrotetrazoliuim chloride solution was added onto each well. After incubating overnight at 37°C, the colonies were captured by an image microscope and quantified after a whole plate scan.

### Cell invasion assay

MDA-MB-231 and MDA-MB-468 cell invasion assay was performed using BD BioCoat Matrigel Invasion Chamber (354480) according to the manufacturer’s instructions. In total, 3 × 10^4^ (for MDA-MB-231) and 3 × 10^5^ (for MDA-MB-468) cells were inoculated into each chamber in triplicate and incubated for 18 h at 37 °C, 5% CO2 incubator. The cells on the lower surface of the membrane were stained using Diff-Quick stain kit (B4132-1A) from SIEMENS, and then counted under EVOS XL Core Microscope (Cat# AMEX1000, Thermo Fisher Scientific).

### RNA-seq Analysis

Procedures was described previously (Liao *et al*, 2020). Briefly, Total RNA from triplicates was extracted from MDA-MB-231 cells infected with control or ZHX2 shRNAs by using RNeasy kit with on column DNase digestion (Qiagen). Library preparation and sequencing were performed by BGI as paired end 50bp reads. Reads were then filtered for adapter contamination using cutadapt (Patro *et al*, 2017) and filtered such that at least 90% of bases of each read had a quality score >20. Reads were aligned to the reference genome (hg19) using STAR version 2.5.2b, and only primary alignments were retained (Love *et al*, 2014). Reads overlapping blacklisted regions of the genome were then removed. Transcript abundance was then estimated using salmon (Miller *et al*, 2012), and differential expression was detected using DESeq2 (Bird *et al*, 2010). RNA-seq data are available at GSE175487. Pathway enrichments were calculated using g:Profiler (Reimand *et al*, 2019) where the pathway database was supplemented with the list of HIF DKO downregulated genes to obtain the adjusted p-value indicating a significant association. Association with the HALLMARK hypoxia pathway was conducted using GSEA (Reimand *et al*., 2019).

### Real-Time PCR

Total RNA was isolated with RNeasy mini kit (Qiagen). First strand cDNA was generated with an iScript cDNA synthesis kit (BioRad). Real-time PCR was performed in triplicate. Real time PCR primer sequences are listed in Table S5, Supplementary.

### ChIP-seq Analyses

MDA-MB-231 cells were infected by HA-ZHX2 which is resistant to ZHX2 sh45, and then infected by ZHX2 sh45. ChIP was performed with HA tag (Cell Signaling, 3724). The ChIP-Seq library was prepared using a ChIP-Seq DNA sample preparation kit (Illumina) according to manufacturer’s instructions. Samples were sequenced on an Illumina HiSeq2500 with single-end 76 bp reads. Reads were then filtered for adaptor contamination using Cutadapt and filtered such that at least 90% of bases of each read had a quality score > 20. Duplicated sequences were then capped at a maximum of 5 occurrences, and reads were aligned to the reference genome (hg19) using STAR (Dobin *et al*, 2013) version 2.5.2b retaining only primary alignments. Reads overlapping blacklisted regions of the genome were then removed. Reads were then extended *in silico* to a fragment size of 250 bp, and regions of significant enrichment relative to input control were identified using MACS2 (Zhang *et al*, 2008). A unified set of enriched regions for ZHX2 was obtained by taking the intersection of the two replicates using bedtools (Quinlan & Hall, 2010). ChIP-seq data for HIF1α was obtained from GSE108833, and data for H3K4me3 and H3K27ac were obtained from GSE49651. ChIP enrichment heatmaps over promoters were generated using deepTools (Ramirez *et al*, 2016).

### Orthotopic Tumor Xenograft

Procedures for animal studies was described previously (Liao *et al*., 2020). Briefly, six-week-old female NOD SCID Gamma mice (NSG, Jackson lab) were used for xenograft studies. Approximately 1×10^6^ viable MDA-MB-231 cells expressing Teton control or Teton ZHX2 shRNAs were resuspended in 1: 1 ratio in 50 μl medium and 50 μl matrigel (Corning, 354234) and injected orthotopically into the fourth mammary fat pad of each mouse. After cell injection and following two consecutive weeks of tumor monitoring to ensure the tumor was successfully implanted, mice were fed Purina rodent chow with doxycycline (Research Diets Inc., #5001). Tumor size was measured twice a week using an electronic caliper. Tumor volumes were calculated with the formula: volume = (L × W^2^)/2, where L is the tumor length and W is the tumor width measured in millimeters. The rough mass of tumors was presented as mean ± SEM and evaluated statistically using *t* test. After mice were sacrificed, lung ex vivo imaging was performed immediately to examine tumor metastasis. All animal experiments were in compliance with National Institutes of Health guidelines and were approved by the University of Texas, Southwestern Medical Center Institutional Animal Care and Use Committee.

### Binding Free Energy Calculations using MM/PBSA

To calculate the binding Gibbs free energy change between two groups of molecules, we used Molecular Mechanics Poisson– Boltzmann Surface Area continuum solvation (MM/PBSA) approach(Miller *et al*., 2012) to analyze trajectories out of Molecular Dynamics (MD) simulations (detailed in Supporting Information). In this approach, the binding Gibbs free energy change, ΔGbind, can be expressed in terms of the xyz (ΔGMM), xyz (ΔGsolv), and the entropy of the system (TΔS) as shown in equation (1).

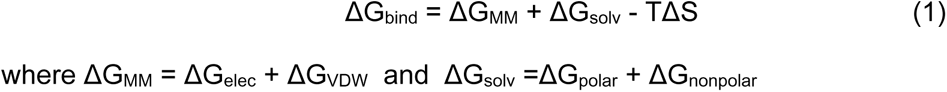

### Statistical Analysis

All statistical analysis was conducted using Prism 8.0 (GraphPad Software). All graphs depict mean ± SEM unless otherwise indicated. Statistical significances are denoted as n.s. (not significant; P>0.05), *P < 0.05, **P < 0.01, ***P < 0.001, ****P < 0.0001. The numbers of experiments are noted in Figure legends. To assess the statistical significance of a difference between two conditions, we used unpaired two-tail student’s *t*-test. For experiments comparing more than two conditions, differences were tested by a one-way ANOVA followed by Dunnett’s or Tukey’s multiple comparison tests.

### Data availability

RNA-Seq and CHIP-Seq data are available GEO175487. All data generated or analysed during this study are included in the manuscript and supporting files.

## Acknowledgments

This work was supported in part by the National Cancer Institute (Q.Z., R01CA211732 and R21CA223675), Cancer Prevention and Research Institute of Texas (CPRIT, RR190058 to Q.Z) and American Cancer Society (RSG-18-059-01-TBE). J.M.S. and T.S.P. were supported by NINDS (P30NS045892). Q.Z is an American Cancer Society Research Scholar, CPRIT Scholar in Cancer Research, V Scholar, Kimmel Scholar, Susan G. Komen Career Catalyst awardee and Mary Kay Foundation awardee.

## Author contributions

C.L. W.F. and Q.Z. conceived, performed and interpreted experiments. W.F. performed the ChIP sequencing experiment. J.M.S., T.S.P. and C.F. performed the RNA-seq and ChIP-seq bioinformatics analyses. C.L. and R.S. performed the animal studies. C.L.O., H.L. performed data-driven analyses and structural simulations. Y.Y., L.H., R.S., U.M., W.L., W.K., W.L. and L.Y. helped to provide critical advice and revisions for the paper. Q.Z., C.L. and W.F. wrote the paper with critical comments from all authors.

## Supplemental Methods

### Survival analysis

The K-M plots were got from https://kmplot.com(9). We chose TNBC patients as follow: ER status-IHC: ER negative, ER status-array: ER negative, PR status-IHC: PR negative, HER2 status-array: HER2 negative. Finally, 153 TNBC patients were included in the overall survival (OS) analysis. ZHX2 overexpression were chosen as upper tertile expression.

### Luciferase reporter assay

For HIF transcription assay, sub-confluent MDA-MB-231 cells (200,000 cells/24-well plate) were transiently transfected with 30 ng pCMV-Renilla and100 ng of HRE-Luci reporter. Forty-eight hours after transfection, luciferase assays were performed by Dual-Luciferase® Reporter Assay System (Promega, E1960). The experiments were repeated in triplicate with similar results.

### DNA-protein contact analysis from structural bioinformatics data

Sequences in the apo-form human homeodomains 2,3 and 4 (HD2/3/4) (PDB ID: 3NAU, 2DMP and 3NAR) (Bird et al., 2010) (Bank, 2020) were BLASTed against the sequences in the Protein Data Bank (PDB)(Berman et al., 2000). Among the resolved structures, 12, 20 and 15 DNA-bound complexes with sequence identity higher than 30% (Rost, 1999) for HD2, HD3, and HD4 were identified. We then carried out the homology-modeling, using SWISS-MODEL (Waterhouse et al., 2018), to structurally model the HD2/3/4 using their corresponding bound-forms as the templates, so that their sequences assume the protein structures in the DNA-complexed forms. For instance, HD2 sequence could therefore adapt 12 different bound-form protein structures. With this method, we found every HD protein contact the DNA with its last (C-terminal) helix (Supplemental Figure 5). We then count the number of DNA-protein contacts at the atomic level for every residue in the C-terminal helix. The top-ranked residues in HD2/3/4, in terms of their DNA contact frequency normalized by the number of bound-forms used, are listed together with their evolutionary conservation in Supplemental Table 1.

### MD simulations

#### Homology Modeling to create DNA-bound HD complexes for simulations

Because there are no experimentally solved DNA-complexed structures for human HD2, 3 and 4, in order to simulate the human HD-DNA interaction, our goal is to find structurally solved DNA-bound forms whose DNA sequence could have the highest chance to stably interact with human HD2/3/4. We aimed to ensure the highest likelihood of stable interaction between the selected DNA and HD2/3/4 as well as to have a fair comparison of binding ability for HD2/3/4 and their DNA-binding residues. To this end, we searched the DNA-bound HD proteins containing a DNA-binding helical stretch that has the highest sequence identity with the C-terminal helices (the main DNA-binding helix; see Supplemental Figure 1 and Video 1) in HD2, 3 and 4, respectively. Among the bound-form proteins that have the top 2 highest sequence homology in the C-terminal helices with those in human HD2/3/4, by homology modeling using SWISS-MODEL (Waterhouse et al., 2018), a NMR-resolved structural ensemble of VND/NK-2 homeodomain-DNA complex (PDB ID: 1NK2, where the first model is taken) (Gruschus, Tsao, Wang, Nirenberg, & Ferretti, 1997) was chosen to build the DNA-bound form of HD2, an X-ray resolved structure of Yeast MATα2 homeodomain/MCM1 transcription factor/DNA complex (PDB ID: 1MNM;) (Tan & Richmond, 1998)was chosen to build the DNA-bound form of HD3, and an X-ray resolved structure of Oct-1 Transcription factor DNA complex (PDB ID:1HF0;) (Remenyi et al., 2001) was chosen to build the DNA-bound form of HD4, respectively. The homology of C-terminal helices between HD2/3/4 and their corresponding bound-form templates are 54.55%, 53.85%, and 100% respectively.

#### System Setup and Energy Minimization

Prior to solvation and addition of ions, protonation state and the net charge of HD2/3/4-dsDNA complexes at pH 7.0 were calculated using PDB2PQR(Dolinsky et al., 2007). The starting structure was prepared using ff14SB (Maier et al., 2015)force-fields for proteins, bsc1 (Ivani et al., 2016) force fields for the DNA, TIP3P water model, and monovalent ion parameters (Joung & Cheatham, 2009) through tLeap(Daoudi et al., 2019) from AmberTools18. To neutralize the charge of each system, 24 Na+ were added into hb2-dsDNA and hb3-dsDNA complexes and 23 Na+ were added into hb4-dsDNA complex systems. In addition, 23 Na^+^ and 23 Cl^-^ ions were added to each system to reach 100 mM salt concentration. Each system was prepared in a water box measuring 78Å on all sides.

Energy minimization for each of the systems was done in two stages. In the first stage, a harmonic restraint of 100 kcal/mol/Å^2^ was applied on all heavy atoms of both protein and dsDNA. In the second stage, the harmonic restraints for protein’s CA atoms were relaxed to 2 kcal/mol/Å^2^ while all the DNA’s heavy atoms were still subject to a 100 kcal/mol/Å^2^ restraint.

#### Equilibration and Explicit Solvent Production MD Simulations

Each energy-minimized system was gradually heated from 50K to 320K and cooled down to 310K in a canonical (NVT) ensemble, using Langevin thermostat (Pastor, Brooks, & Szabo, 1988) with a collision frequency of 2 ps^-1^, for 25 ps while applying harmonic restraints of 10 kcal/mol/Å^2^ on dsDNA’s C2, C4’, and P atoms and 2 kcal/mol/Å^2^ on protein’s CA atoms. Each of the systems was equilibrated first in a canonical ensemble at 310K for 15 ns. This was followed by an isothermal-isobaric ensemble for 20 ns at 310K applying harmonic restraints of 2 kcal/mol/Å^2^ on dsDNA’s C2, C4’, and P atoms and 1 kcal/mol/Å^2^ on protein’s CA atoms. Further equilibration isothermal-isobaric ensemble (NPT), where a constant pressure was maintained by Berendsen barostat (H. J. C. Berendsen, 1998) at 1 atm and 310K, was done for 40 ns, while harmonic restraints of 2 kcal/mol/Å^2^ on dsDNA’s C2, C4’, and P atoms and 0.1 kcal/mol/Å^2^ on protein’s CA atoms were applied. This was followed by a 350 ns production run at 2 fs time step applying the SHAKE constraint algorithm (Hopkins, Le Grand, Walker, & Roitberg, 2015) to hydrogen atoms in isothermal-isobaric ensemble at 310K and 1 atm. All the simulations were carried out by the AMBER18 software package (Daoudi et al., 2019) with long-range electrostatic forces being calculated using Particle Mesh Ewald method (Tom Darden, 1993) at a 10Å cutoff distance.

## Legend for Figure-source data

**Figure 1-source data**. Uncropped western blot images for Figure 1.

**Figure 2-source data**. Uncropped western blot images for Figure 2.

**Figure 3-source data**. Uncropped western blot images for Figure 3.

**Figure 4-source data**. Uncropped western blot images for Figure 4.

**Figure 6-source data**. Uncropped western blot images for Figure 6.

**Figure supplement 1-source data**. Uncropped western blot images for Figure supplement 1.

**Figure supplement 2-source data**. Uncropped western blot images for Figure supplement 2.

**Figure supplement 3-source data**. Uncropped western blot images for Figure supplement 3.

**Figure supplement 4-source data**. Uncropped western blot images for Figure supplement 4.

## Supplemental Figures

**Figure S1.**
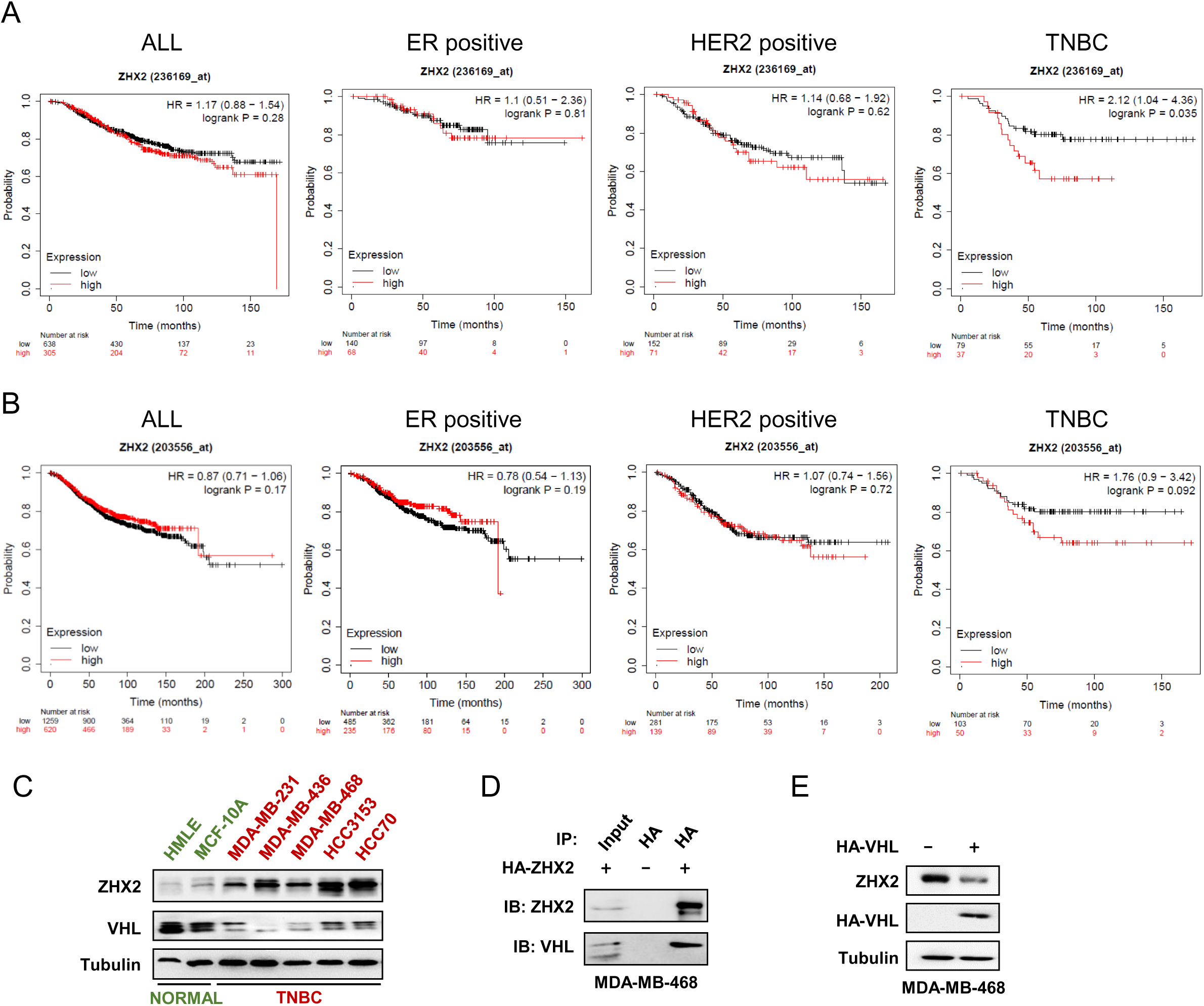
ZHX2 overexpression leads worse survival and is potentially regulated by pVHL in breast cancer. (**A-B**) The overall survival of breast cancer patients with high or low ZHX2 expression in different breast cancer subtypes. The K-M plots were generated from https://kmplot.com using two Affymetrix probe, ZHX2: 236169_at (**A**) and 203556_at (**B**). (**C**) Immunoblots of lysates from normal breast epithelial cell and TNBC cell lines. (**D**) Immunoprecipitations of MDA-MB-468 cells expressing either control vector or HA-ZHX2. (**E**) Immunoblots of lysates from MDA-MB-468 cells infected with either control vector or HA-VHL.

**Figure S2.**
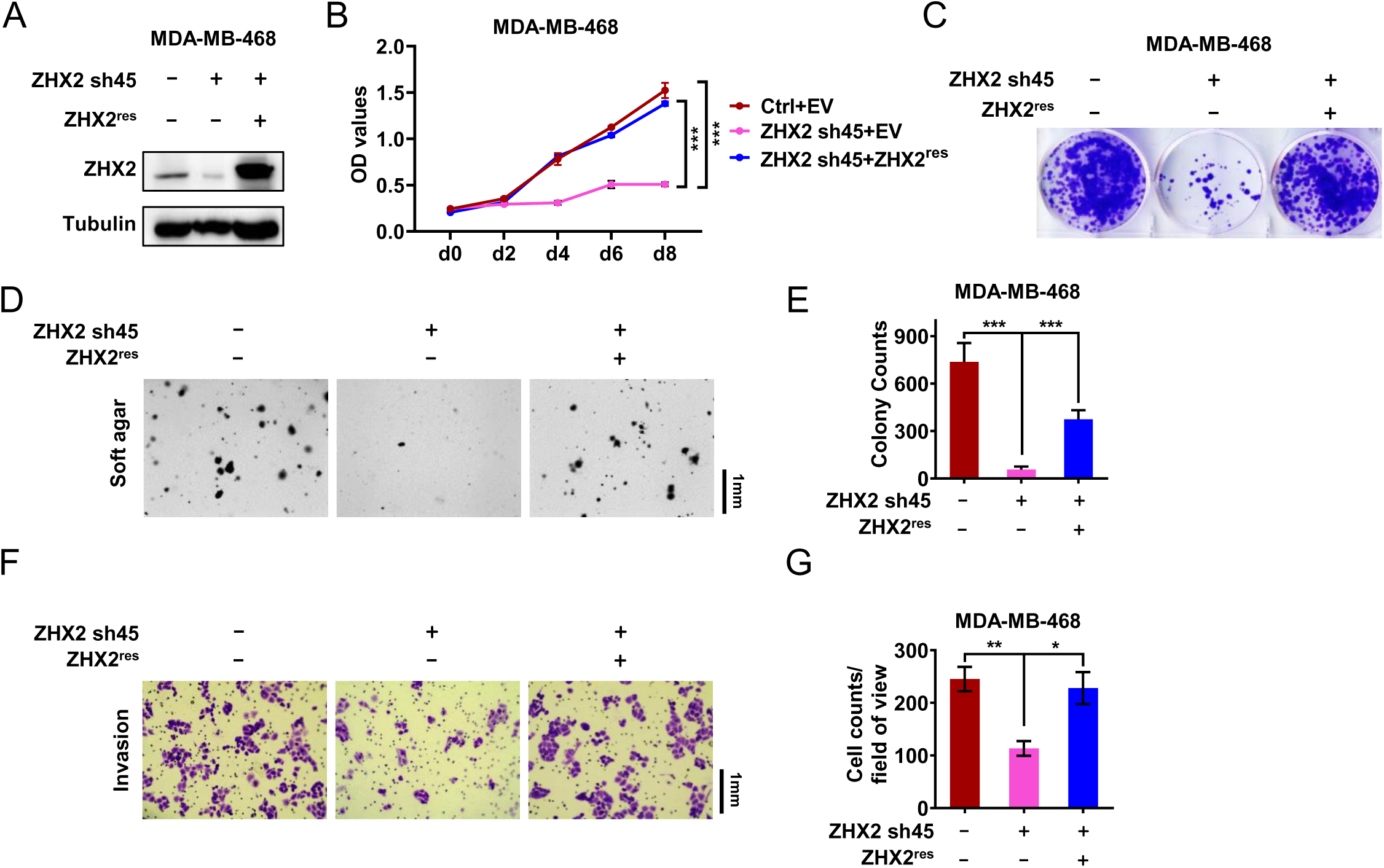
The phenotype of ZHX2 shRNA on cell proliferation and invasion is due to its on-target effect. (**A**) Immunoblot of cell lysates of MDA-MB-468 cells transfected with ZHX2 sh45-resistant ZHX2 (ZHX2^res^) or empty (EV) vector, followed by ZHX2 sh45 or control (Ctrl) shRNA infection. (**B**) Cell proliferation assays of MDA-MB-468 cells transfected with sh45-resistant ZHX2 (ZHX2^res^) or empty (EV) vector, followed by ZHX2 sh45 or control (Ctrl) shRNA infection. (**C**) 2-D colony formation assay of MDA-MB-468 cells transfected with sh45-resistant ZHX2 (ZHX2^res^) or empty (EV) vector, followed by ZHX2 sh45 or control (Ctrl) shRNA infection. (**D-E**) Representative soft agar colony (**D**) and quantification (**E**) of MDA-MB-468 cells transfected with ZHX2 sh45-resistant ZHX2 (ZHX2^res^) or empty (EV) vector, followed by ZHX2 sh45 or control (Ctrl) shRNA infection. (**F-G**) Invasion assays (**F**) and quantification (**G**) of MDA-MB-468 cells transfected with ZHX2 sh45-resistant ZHX2 (ZHX2^res^) or empty (EV) vector, followed by ZHX2 sh45 or control (Ctrl) shRNA infection.

**Figure S3.**
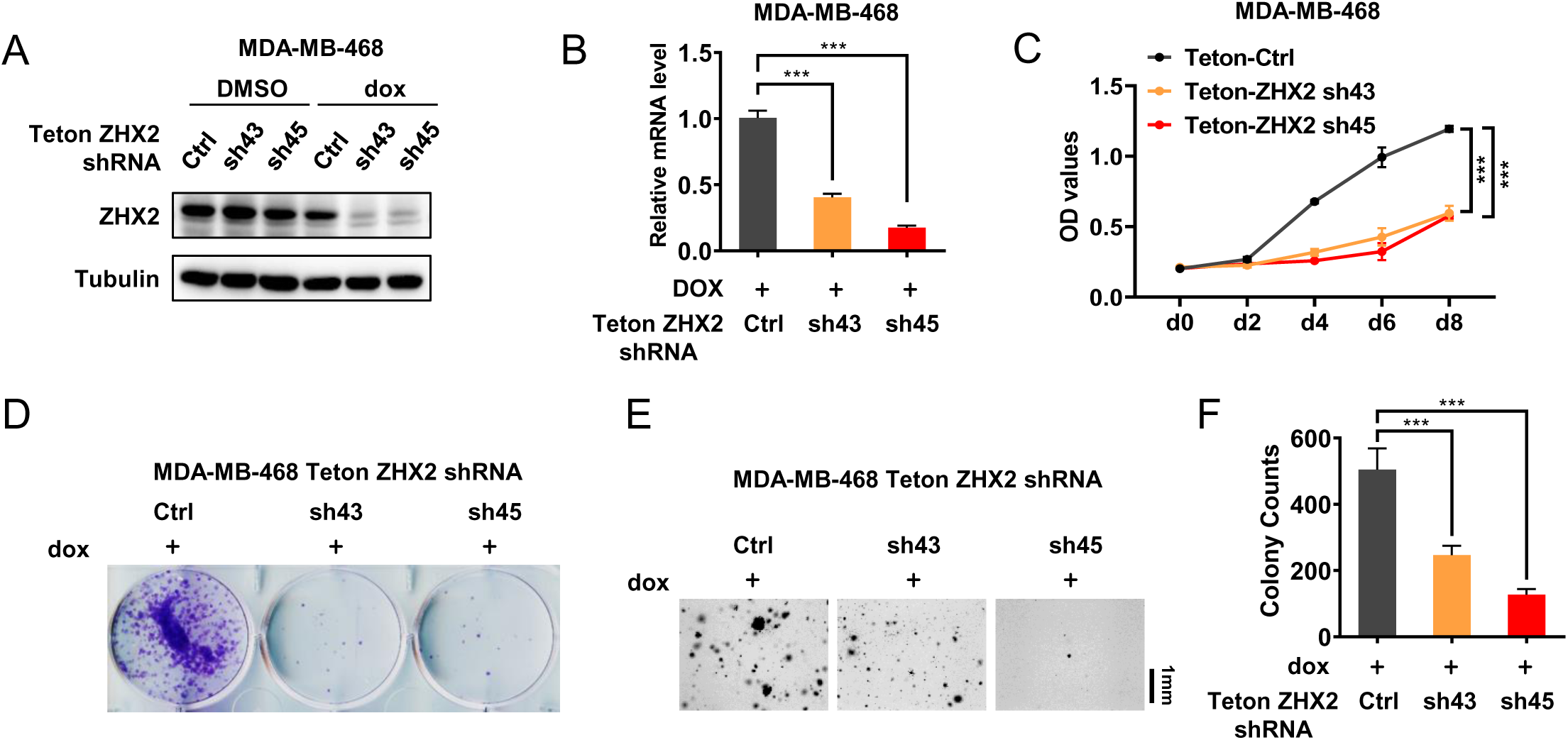
ZHX2 is important for maintaining TNBC tumorigenesis *in vivo*. **(A-B**) Immunoblot (**A**) and qRT-PCR (**B**) of MDA-MB-468 cells infected with lentivirus encoding either Teton-ZHX2 shRNA (43, 45) or Teton-Ctrl. (**C-D**) Cell proliferation assay (**C**) and 2-D colony formation assay (**D**) of MDA-MB-468 cells infected with lentivirus encoding either Teton-ZHX2 shRNA (43, 45) or Teton-Ctrl. (**E-F**) Representative soft agar colony (**E**) and quantification (**F**) of MDA-MB-468 cell lines infected with lentivirus encoding either Teton-ZHX2 shRNA (43, 45) or Teton-Ctrl.

**Figure S4.**
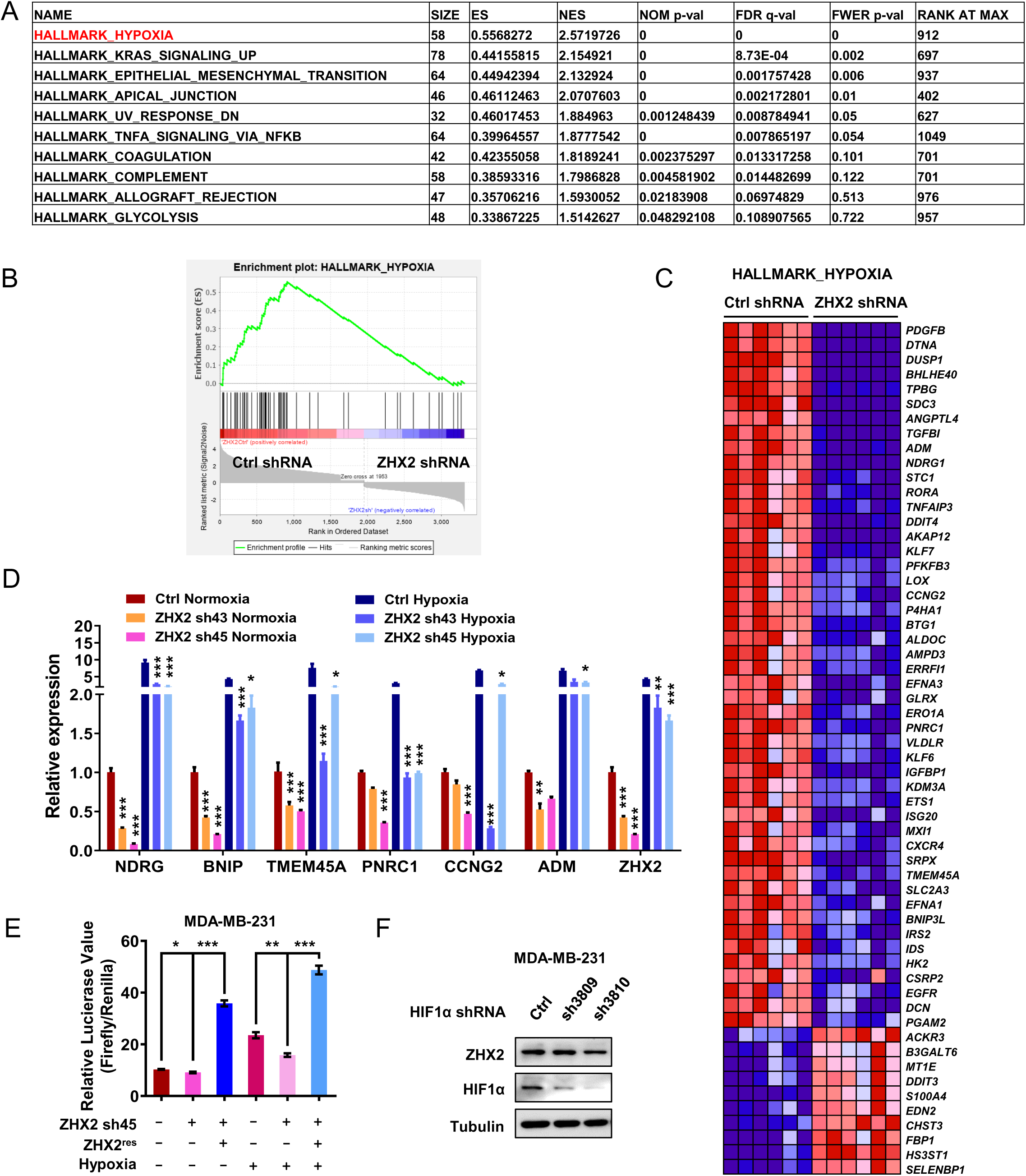
ZHX2 regulates HIF1 signaling in TNBC. (**A**) Gene set enrichment analysis (GSEA) of the significantly decreased pathways in ZHX2 depleted MDA-MB-231 cells. (**B-C**) GSEA plot (**B**) and gene set heatmap (**C**) suggest hypoxia pathway is significantly downregulated in ZHX2 depleted MDA-MB-231 cells. (**D**) qRT-PCR quantification of relative mRNA expression of hypoxia target genes from MDA-MB-231 cells infected with ZHX2 shRNA 43, 45 or Ctrl under normoxia or hypoxia conditions. (**E**) HRE double luciferase gene assay of MDA-MB-231 cells infected with ZHX2 sh45, sh45-resistant ZHX2 (ZHX2^res^) or Ctrl under normoxia or hypoxia condition. (**F**) Immunoblot of MDA-MB 231 cells infected with lentivirus encoding either HIF1α shRNA 3809, 3810 or Ctrl.

**Figure S5.**
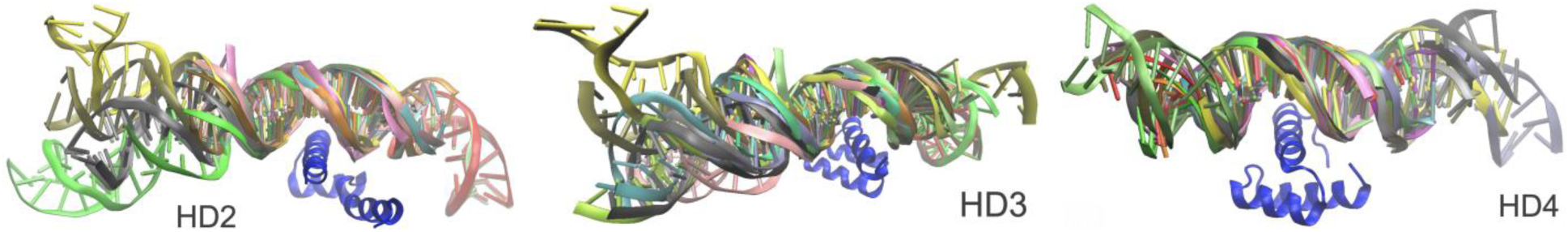
Residue-based DNA contact analysis derived from ensembles of DNA-bound HD proteins.

## Supplemental Tables

**Table S1.** Amplification status of ZHX2 in different breast cancer subtype.

| Dataset | Breast cancer patients |  |  |  | TNBC patients |  |  |  | ER positive |  |  |  | HER2 positive |  |  |  |
| --- | --- | --- | --- | --- | --- | --- | --- | --- | --- | --- | --- | --- | --- | --- | --- | --- |
|  | Patient | Sample | ZHX2 Amp | Percentage | Patient | Sample | ZHX2 Amp | Percentage | Patient | Sample | ZHX2 Amp | Percentage | Patient | Sample | ZHX2 Amp | Percentage |
| Breast Cancer (METABRIC, Nature 2012 & Nat Commun 2016) | 2509 | 2509 | 512 | 23.6 | 320 | 320 | 104 | 32.5 | 1825 | 1825 | 341 | 21.1 | 247 | 247 | 81 | 32.8 |
| The Metastatic Breast Cancer Project (Provisional, February 2020) | 180 | 237 | 44 | 18.6 | 5 | 6 | 2 | 33.3 | 94 | 126 | 18 | 14.3 | 37 | 44 | 10 | 22.7 |
| Breast Invasive Carcinoma (TCGA, Cell 2015) | 817 | 818 | 148 | 18.1 | 83 | 83 | 25 | 30.1 | 601 | 601 | 90 | 15.0 | 121 | 121 | 31 | 25.6 |
| Breast Invasive Carcinoma (TCGA, Firehose Legacy) | 1101 | 1108 | 197 | 18.2 | 116 | 117 | 36 | 32.1 | 808 | 814 | 120 | 15.1 | 164 | 164 | 39 | 24.1 |
| Breast Invasive Carcinoma (TCGA, Nature 2012) | 825 | 825 | 92 | 11.8 | 123 | 123 | 23 | 19.3 | 601 | 601 | 58 | 10.3 | 114 | 114 | 20 | 18.2 |

**Table S2.** Top DNA-contacting residues in HD2/3/4 along with their evolutionary conservation.

| Homeobox 2 |  |  | Homeobox 3 |  |  | Homeobox 4 |  |  |
| --- | --- | --- | --- | --- | --- | --- | --- | --- |
| Residue | Mean Total Contacts | Conservation Score* | Residue | Mean Total Contacts | Conservation Score* | Residue | Mean Total Contacts | Conservation Score* |
| LYS 485 | 52.50 | 5 | **GLU 579** | 31.05 | 8 | **ARG 674** | 129.20 | 6 |
| **ARG 491** | 36.33 | 9 | **ARG 581** | 24.80 | 6 | **GLU 678** | 58.73 | 8 |
| PHE 463 | 32.92 | 2 | SER 575 | 23.20 | 5 | GLU 671 | 45.80 | 7 |
| ARG 493 | 25.08 | 5 | **LYS 582** | 23.00 | 7 | **ARG 680** | 28.33 | 9 |
| **ASP 489** | 22.42 | 8 | PHE 553 | 13.00 | 1 | LYS 635 | 27.73 | 9 |
| GLU 482 | 14.25 | 6 | GLU 572 | 9.55 | 8 | LYS 684 | 25.13 | 7 |
| LYS 484 | 11.42 | 7 | ARG 570 | 9.25 | 7 | LYS 677 | 23.47 | 7 |
| TYR 492 | 9.58 | 5 | TRP 576 | 7.55 | 9 | TRP 652 | 15.53 | 4 |
| ARG 480 | 8.67 | 8 | ASP 585 | 4.70 | 7 | TRP 675 | 10.13 | 9 |
| TRP 486 | 7.42 | 9 | SER 578 | 4.70 | 5 | CYS 681 | 6.33 | 7 |

\*obtained using ConSurf server (9 – most conserved, 1 – most diverse); bold-faced residues were chosen for the mutagenesis study on TNBC suppression. DNA-protein contact is defined by heavy atom contact within 4Å, while the same DNA atoms contacted by different protein atoms are considered as separate contacts.

**Table S3.** MM/PBSA Calculations of the Free Energy (ΔH-TΔS) for Homeobox 2-4 dsDNA complexes.

| | $\Delta H$ (kcal/mol) | $(T\Delta S)$ (kcal/mol) | $\Delta G$ (kcal/mol) |
| --- | --- | --- | --- |
| **HD2+DNA** | -60.25 ± 0.50 | -48.05 ± 0.97 | **-12.20 ± 1.09** |
| **HD3+DNA** | -56.93 ± 0.46 | -45.53 ± 2.39 | **-11.40 ± 2.43** |
| **HD4+DNA** | -39.00 ± 0.50 | -31.23 ± 2.34 | **-7.77 ± 2.39** |

**Table S4.** Top C-terminal helix residues in the Homeobox 2-4 contributing the most DNA binding enthalpy.

| Homeobox 2 |  | Homeobox 3 |  | Homeobox 4 |  |
| --- | --- | --- | --- | --- | --- |
| Residue number** | $\Delta H_{\text{total}}$ (kcal/mol)* | Residue number** | $\Delta H_{\text{total}}$ (kcal/mol)* | Residue number** | $\Delta H_{\text{total}}$ (kcal/mol)* |
| ARG 491 | $-8.75 \pm 0.17$ | ARG 581 | $-12.51 \pm 0.15$ | ARG 674 | $-6.81 \pm 0.12$ |
| ARG 480 | $-7.98 \pm 0.08$ | ARG 571 | $-7.54 \pm 0.09$ | LYS 684 | $-3.46 \pm 0.08$ |
| ARG 493 | $-7.41 \pm 0.09$ | LYS 582 | $-6.98 \pm 0.10$ | LYS 677 | $-3.18 \pm 0.07$ |
| LYS 484 | $-6.49 \pm 0.08$ | ARG 580 | $-4.43 \pm 0.02$ | ARG 680 | $-2.41 \pm 0.01$ |
| LYS 485 | $-5.00 \pm 0.11$ | ARG 570 | $-2.66 \pm 0.02$ | ARG 669 | $-1.78 \pm 0.01$ |
| TYR 492 | $-2.88 \pm 0.03$ | ARG 584 | $-2.24 \pm 0.01$ | TRP 675 | $-1.45 \pm 0.05$ |
| ARG 496 | $-2.12 \pm 0.11$ | SER 578 | $-0.71 \pm 0.06$ | LEU 682 | $-0.79 \pm 0.02$ |
| GLN 495 | $-1.86 \pm 0.07$ | PHE 577 | $-0.35 \pm 0.01$ | CYS 681 | $-0.36 \pm 0.02$ |
| TRP 486 | $-0.96 \pm 0.06$ | LEU 583 | $-0.22 \pm 0.00$ | THR 670 | $-0.36 \pm 0.04$ |
| HID 490 | $-0.64 \pm 0.01$ | | | ASN 679 | $-0.24 \pm 0.02$ |
| SER 481 | $-0.49 \pm 0.03$ | | | | |
| PHE 487 | $-0.28 \pm 0.01$ | | | | |
| CYS 494 | $-0.21 \pm 0.00$ | | | | |

\* $\Delta H_{\text{total}}$ : enthalpic binding energy derived from MM/PBSA, \*\*this numbering is based on the Uniprot database where the sequence was obtained, highlighted in blue represents the mutated amino acid in the experimental results

**Table S5.**
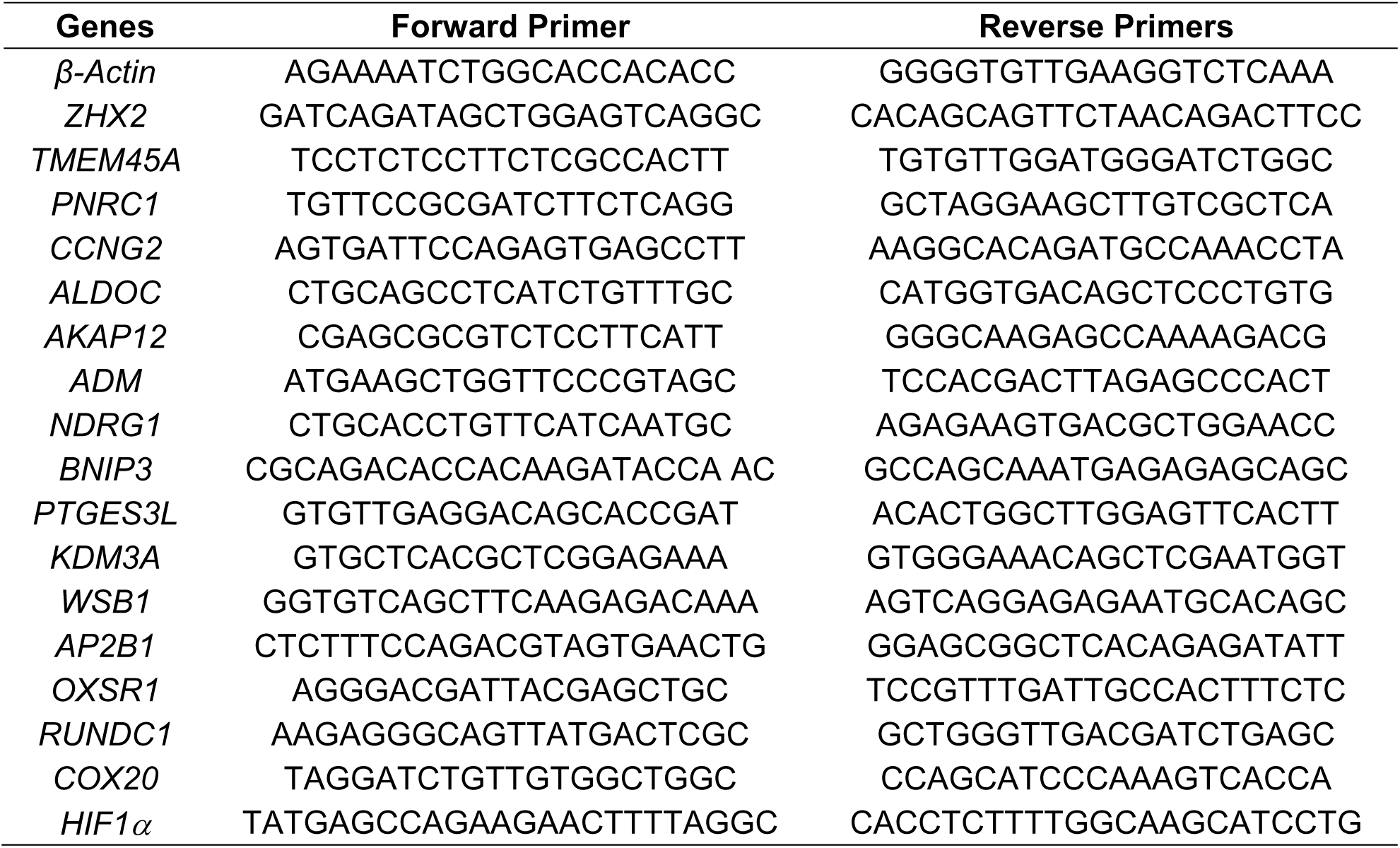
Real-time PCR primers used in this study.

**Movie S1.**

The 350-ns MD simulation trajectory of 1NK2-based HB2-dsDNA complex. The HB2-dsDNA complex is in secondary structure cartoon representation. The top C-terminal helix residue that most contacts the DNA, which is ARG491, and the DNA atoms within 4Å distance from it are in licorice representation.

**Movie S2.**

The 350-ns MD simulation trajectory of 1MNM-based HB3-dsDNA complex. The HB3-dsDNA complex is in secondary structure cartoon representation (violet color: HB3’s helix; grey color: dsDNA). The top C-terminal helix residue that most contacts the DNA, which is ARG581, and the DNA atoms within 4Å distance from it are in licorice representation.

**Movie S3.**

The 350-ns MD simulation trajectory of 1HF0-based HB4-dsDNA complex. The HB4-dsDNA complex is in secondary structure cartoon representation (violet color: HB4’s helix; grey color: dsDNA). The top C-terminal helix residue that most contacts the DNA, which is ARG674, and the DNA atoms within 4Å distance from it are in licorice representation.

